# Systematic integration of biomedical knowledge prioritizes drugs for repurposing

**DOI:** 10.1101/087619

**Authors:** Daniel S. Himmelstein, Antoine Lizee, Christine Hessler, Leo Brueggeman, Sabrina L. Chen, Dexter Hadley, Ari Green, Pouya Khankhanian, Sergio E. Baranzini

## Abstract

The ability to computationally predict whether a compound treats a disease would improve the economy and success rate of drug approval. This study describes Project Rephetio to systematically model drug efficacy based on 755 existing treatments. First, we constructed Hetionet (neo4j.het.io), an integrative network encoding knowledge from millions of biomedical studies. Hetionet v1.0 consists of 47,031 nodes of 11 types and 2,250,197 relationships of 24 types. Data was integrated from 29 public resources to connect compounds, diseases, genes, anatomies, pathways, biological processes, molecular functions, cellular components, pharmacologic classes, side effects, and symptoms. Next, we identified network patterns that distinguish treatments from non-treatments. Then we predicted the probability of treatment for 209,168 compound–disease pairs (het.io/repurpose). Our predictions validated on two external sets of treatment and provided pharmacological insights on epilepsy, suggesting they will help prioritize drug repurposing candidates. This study was entirely open and received realtime feedback from 40 community members.

## Introduction

The cost of developing a new therapeutic drug has been estimated at 1.4 billion dollars [1], the process typically takes 15 years from lead compound to market [2], and the likelihood of success is stunningly low [3]. Strikingly, the costs have been doubling every 9 years since 1970, a sort of inverse Moore’s law, which is far from an optimal strategy from both a business and public health perspective [4]. Drug repurposing — identifying novel uses for existing therapeutics — can drastically reduce the duration, failure rates, and costs of approval [5]. These benefits stem from the rich preexisting information on approved drugs, including extensive toxicology profiling performed during development, preclinical models, clinical trials, and postmarketing surveillance.

Drug repurposing is poised to become more efficient as mining of electronic health records (EHRs) to retrospectively assess the effect of drugs gains feasibility [6–9]. However, systematic approaches to repurpose drugs based on mining EHRs alone will likely lack power due to multiple testing. Similar to the approach followed to increase the power of genome-wide association studies (GWAS) [10, 11], integration of biological knowledge to prioritize drug repurposing will help overcome limited EHR sample size and data quality.

In addition to repurposing, several other paradigm shifts in drug development have been proposed to improve efficiency. Since small molecules tend to bind to many targets, polypharmacology aims to find synergy in the multiple effects of a drug [12]. Network pharmacology assumes diseases consist of a multitude of molecular alterations resulting in a robust disease state. Network pharmacology seeks to uncover multiple points of intervention into a specific pathophysiological state that together rehabilitate an otherwise resilient disease process [13,14]. Although target centric drug discovery has dominated the field for decades, phenotypic screens have more recently resulted in a comparatively higher number of first-in-class small molecules [15]. Recent technological advances have enabled a new paradigm in which mid-to high-throughput assessment of intermediate phenotypes, such as the molecular response to drugs, is replacing the classic target discovery approach [16–18]. Furthermore, integration of multiple channels of evidence, particularly diverse types of data, can overcome the limitations and weak performance inherent to data of a single domain [19]. Modern computational approaches offer a convenient platform to tie these developments together as the reduced cost and increased velocity of *in silico* experimentation massively lowers the barriers to entry and price of failure [20, 21].

Hetnets (short for heterogeneous networks) are networks with multiple types of nodes and relationships. They offer an intuitive, versatile, and powerful structure for data integration by aggregating graphs for each relationship type onto common nodes. In this study, we developed a hetnet (Hetionet v1.0) by integrating knowledge and experimental findings from decades of biomedical research spanning millions of publications. We adapted an algorithm originally developed for social network analysis and applied it to Hetionet v1.0 to identify patterns of efficacy and predict new uses for drugs. The algorithm performs edge prediction through a machine learning framework that accommodates the breadth and depth of information contained in Hetionet v1.0 [22,23]. Our approach represents an *in silico* implementation of network pharmacology that natively incorporates polypharmacology and high-throughput phenotypic screening.

One fundamental characteristic of our method is that it learns and evaluates itself on existing medical indications (i.e. a “gold standard”). Next, we introduce previous approaches that also performed comprehensive evaluation on existing treatments. A 2011 study, named PREDICT, compiled 1,933 treatments between 593 drugs and 313 diseases [24]. Starting from the premise that similar drugs treat similar diseases, PREDICT trained a classifier that incorporates 5 types of drug-drug and 2 types of disease-disease similarity. A 2014 study compiled 890 treatments between 152 drugs and 145 diseases with transcriptional signatures [25]. The authors found that compounds triggering an opposing transcriptional response to the disease were more likely to be treatments, although this effect was weak and limited to cancers. A 2016 study compiled 402 treatments between 238 drugs and 78 diseases and used a single proximity score — the average shortest path distance between a drug’s targets and disease’s associated proteins on the interactome — as a classifier [26].

We build on these successes by creating a framework for incorporating the effects of any biological relationship into the prediction of whether a drug treats a disease. By doing this, we were able to capture a multitude of effects that have been suggested as influential for drug repurposing including drug-drug similarity [24,27], disease-disease similarity [24,28], transcriptional signatures [17,18,25,29,30], protein interactions [26], genetic association [31,32], drug side effects [33,34], disease symptoms [35], and molecular pathways [36]. Our ability to create such an integrative model of drug efficacy relies on the hetnet data structure to unite diverse information. On Hetionet v1.0, our algorithm learns which types of compound–disease paths discriminate treatments from non-treatments in order to predict the probability that a compound treats a disease.

We refer to this study as Project Rephetio (pronounced as **rep**-*het*-*ee*-oh). Both Rephetio and Hetionet are portmanteaus combining the words **rep**urpose, **het**erogeneous, and **net**work with the URL het.**io**.

## Results

### Hetionet v1.0

We obtained and integrated data from 29 publicly available resources to create Hetionet v1.0 (Figure 1). The hetnet contains 47,031 nodes of 11 types (Table 1) and 2,250,197 relationships of 24 types (Table 2). The nodes consist of 1,552 small molecule compounds and 137 complex diseases, as well as genes, anatomies, pathways, biological processes, molecular functions, cellular components, perturbations, pharmacologic classes, drug side effects, and disease symptoms. The edges represent relationships between these nodes and encompass the collective knowledge produced by millions of studies over the last half century.

**Figure 1:**
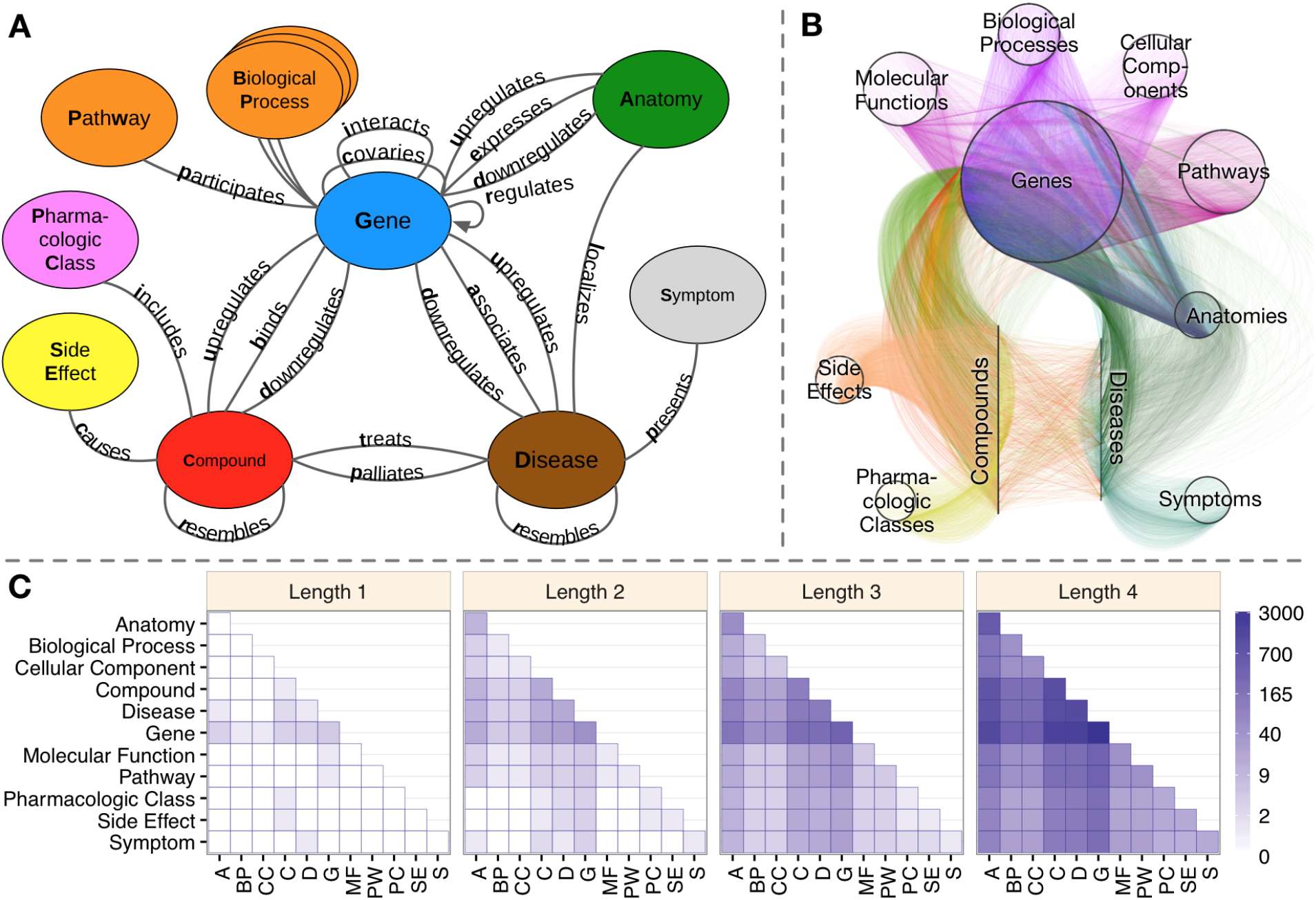
Hetionet v1.0. A) The metagraph, a schema of the network types. B) The hetnet visualized. Nodes are drawn as dots and laid out orbitally, thus forming circles. Edges are colored by type. C) Metapath counts by path length. The number of different types of paths of a given length that connect two node types is shown. For example, the top-left tile in the Length 1 panel denotes that Anatomy nodes are not connected to themselves (i.e. no edges connect nodes of this type between themselves). However, the bottom-left tile of the Length 4 panel denotes that 88 types of length-four paths connect Symptom to Anatomy nodes.

**Table 1:** Metanodes. Hetionet v1.0 includes 11 node types (metanodes). For each metanode, this table shows the abbreviation, number of nodes, number of nodes without any edges, and the number of metaedges connecting the metanode.

| Metanode | Abbr | Nodes | Disconnected | Metaedges |
| --- | --- | --- | --- | --- |
| Anatomy | A | 402 | 2 | 4 |
| Biological Process | BP | 11,381 | 0 | 1 |
| Cellular Component | CC | 1,391 | 0 | 1 |
| Compound | C | 1,552 | 14 | 8 |
| Disease | D | 137 | 1 | 8 |
| Gene | G | 20,945 | 1,800 | 16 |
| Molecular Function | MF | 2,884 | 0 | 1 |
| Pathway | PW | 1,822 | 0 | 1 |
| Pharmacologic Class | PC | 345 | 0 | 1 |
| Side Effect | SE | 5,734 | 33 | 1 |
| Symptom | S | 438 | 23 | 1 |

**Table 2:**
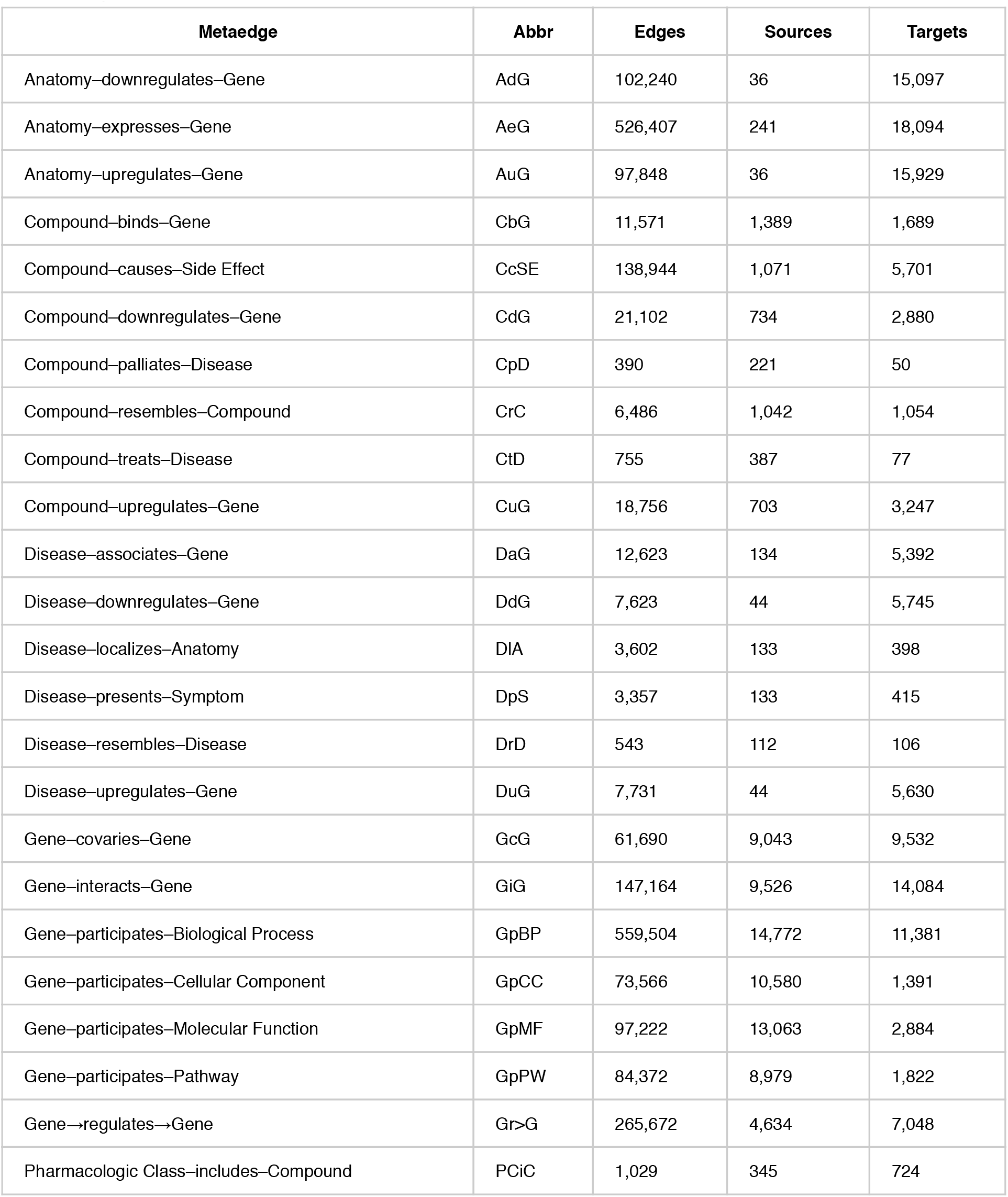
Metaedges. Hetionet v1.0 contains 24 edge types (metaedges). For each metaedge, the table reports the abbreviation, the number of edges, the number of source nodes connected by the edges, and the number of target nodes connected by the edges. Note that all metaedges besides *Gene→regulates→Gene* are undirected.

| Metaedge | Abbr | Edges | Sources | Targets |
| --- | --- | --- | --- | --- |
| Anatomy–downregulates–Gene | AdG | 102,240 | 36 | 15,097 |
| Anatomy–expresses–Gene | AeG | 526,407 | 241 | 18,094 |
| Anatomy–upregulates–Gene | AuG | 97,848 | 36 | 15,929 |
| Compound–binds–Gene | CbG | 11,571 | 1,389 | 1,689 |
| Compound–causes–Side Effect | CcSE | 138,944 | 1,071 | 5,701 |
| Compound–downregulates–Gene | CdG | 21,102 | 734 | 2,880 |
| Compound–palliates–Disease | CpD | 390 | 221 | 50 |
| Compound–resembles–Compound | CrC | 6,486 | 1,042 | 1,054 |
| Compound–treats–Disease | CtD | 755 | 387 | 77 |
| Compound–upregulates–Gene | CuG | 18,756 | 703 | 3,247 |
| Disease–associates–Gene | DaG | 12,623 | 134 | 5,392 |
| Disease–downregulates–Gene | DdG | 7,623 | 44 | 5,745 |
| Disease–localizes–Anatomy | DIA | 3,602 | 133 | 398 |
| Disease–presents–Symptom | DpS | 3,357 | 133 | 415 |
| Disease–resembles–Disease | DrD | 543 | 112 | 106 |
| Disease–upregulates–Gene | DuG | 7,731 | 44 | 5,630 |
| Gene–covaries–Gene | GcG | 61,690 | 9,043 | 9,532 |
| Gene–interacts–Gene | GiG | 147,164 | 9,526 | 14,084 |
| Gene–participates–Biological Process | GpBP | 559,504 | 14,772 | 11,381 |
| Gene–participates–Cellular Component | GpCC | 73,566 | 10,580 | 1,391 |
| Gene–participates–Molecular Function | GpMF | 97,222 | 13,063 | 2,884 |
| Gene–participates–Pathway | GpPW | 84,372 | 8,979 | 1,822 |
| Gene→regulates→Gene | Gr>G | 265,672 | 4,634 | 7,048 |
| Pharmacologic Class–includes–Compound | PCiC | 1,029 | 345 | 724 |

For example, *Compound–binds–Gene* edges represent when a compound binds to a protein encoded by a gene. This information has been extracted from the literature by human curators and compiled into databases such as DrugBank, ChEMBL, DrugCentral, and BindingDB. We combined these databases to create 11,571 binding edges between 1,389 compounds and 1,689 genes. These edges were compiled from 10,646 distinct publications, which Hetionet binding edges reference as an attribute. Binding edges represent a comprehensive catalog constructed from low throughput experimentation. However, we also integrated findings from high throughput technologies — many of which have only recently become available. For example, we generated consensus transcriptional signatures for compounds in LINCS L1000 and diseases in STARGEO.

While Hetionet v1.0 is ideally suited for drug repurposing, the network has broader biological applicability. For example, we have prototyped queries for a) identifying drugs that target a specific pathway, b) identifying biological processes involved in a specific disease, c) identifying the drug targets responsible for causing a specific side effect, and d) identifying anatomies with transcriptional relevance for a specific disease [37]. Each of these queries was simple to write and took less than a second to run on our publicly available Hetionet Browser. While it is possible that existing services provide much of the aforementioned functionality, they offer less versatility. Hetionet differentiates itself in its ability to flexibly query across multiple domains of information. As a proof of concept, we enhanced the biological process query (b), which identified processes that were enriched for disease-associated genes, using multiple sclerosis (MS) as an example disease. The verbose Cypher code for this query is shown below:

~~~
// Specify the type of path to match
(n0:Disease)-[e1:ASSOCIATES_DaG]-(n1:gene)-[:INTERACTES_GIG]-
(n2:gene)-[:PARTICIPATES_GpBp]-(n1:gene)-(n3:BiologicalProcess)
~~~

~~~
// Specify the source and target nodes
n0.name = ‘multiple sclerosis’ AND
n3.name = ‘retina layer formation’
// Require GWAS support for the Disease-associates-Gene relationship
AND ‘GWAS Catalog’ in e1.sources
// Require the interacting gene to be upregulated in a relevant tissue
AND exists((n0)-[:LOCALIZES_DlA]-(:Anatomy)-[:UPREGULATES_AuG]-(n2))
~~~

~~~
RETURN path
~~~

The query above identifies genes that interact with MS GWAS-genes. However, interacting genes are discarded unless they are upregulated in an MS-related anatomy (i.e. anatomical structure, e.g. organ or tissue). Then relevant biological processes are identified. Thus, this single query spans 4 node and 5 relationship types.

The integrative potential of Hetionet v1.0 is reflected by its connectivity. Among the 11 metanodes, there are 66 possible source–target pairs. However, only 11 of them have at least one direct connection. In contrast, for paths of length 2, 50 pairs have connectivity (paths types that start on the source node type and end on the target node type, see Figure 1C). At length 3, all 66 pairs are connected. At length 4, the source–target pair with the fewest types of connectivity (Side Effect to Symptom) has 13 metapaths, while the pair with the most connectivity types (Gene to Gene) has 3,542 pairs. This high level of connectivity across a diversity of biomedical entities forms the foundation for automated translation of knowledge into biomedical insight.

Hetionet v1.0 is accessible via a Neo4j Browser at https://neo4j.het.io. This public Neo4j instance provides users an installation-free method to query and visualize the network. The Browser contains a tutorial guide as well as guides with the details of each Project Rephetio prediction. Hetionet v1.0 is also available for download in JSON, Neo4j, and TSV formats [38]. The JSON and Neo4j database formats include node and edge properties — such as URLs, source and license information, and confidence scores — and are thus recommended.

### Systematic mechanisms of efficacy

One aim of Project Rephetio was to systematically evaluate how drugs exert their therapeutic potential. To address this question, we compiled a gold standard of 755 disease-modifying indications, which form the *Compound–treats–Disease* edges in Hetionet v1.0. Next, we identified types of paths (metapaths) that occurred more frequently between treatments than non-treatments (any compound–disease pair that is not a treatment). The advantage of this approach is that metapaths naturally correspond to mechanisms of pharmacological efficacy. For example, the*Compound–binds–Gene–associates–Disease* (CbGaD) metapath identifies when a drug binds to a protein corresponding to a gene involved in the disease.

We evaluated all 1,206 metapaths that traverse from compound to disease and have length of 2–4 (Figure 2A). To control for the different degrees of nodes, we used the degree-weighted path count (*DWPC*, see Methods) — which downweights paths going through highly-connected nodes [22] — to assess path prevalence. In addition, we compared the performance of each metapath to a baseline computed from permuted networks. Hetnet permutation preserves node degree while eliminating edge specificity, allowing us to isolate the portion of unpermuted metapath performance resulting from actual network paths. We refer to the permutation-adjusted performance measure as Δ AUROC. A positive Δ AUROC indicates that paths of the given type tended to occur more frequently between treatments than non-treatments, after accounting for different levels of connectivity (node degrees) in the hetnet. In general terms, Δ AUROC assesses whether paths of a given type were informative of drug efficacy.

**Figure 2:**
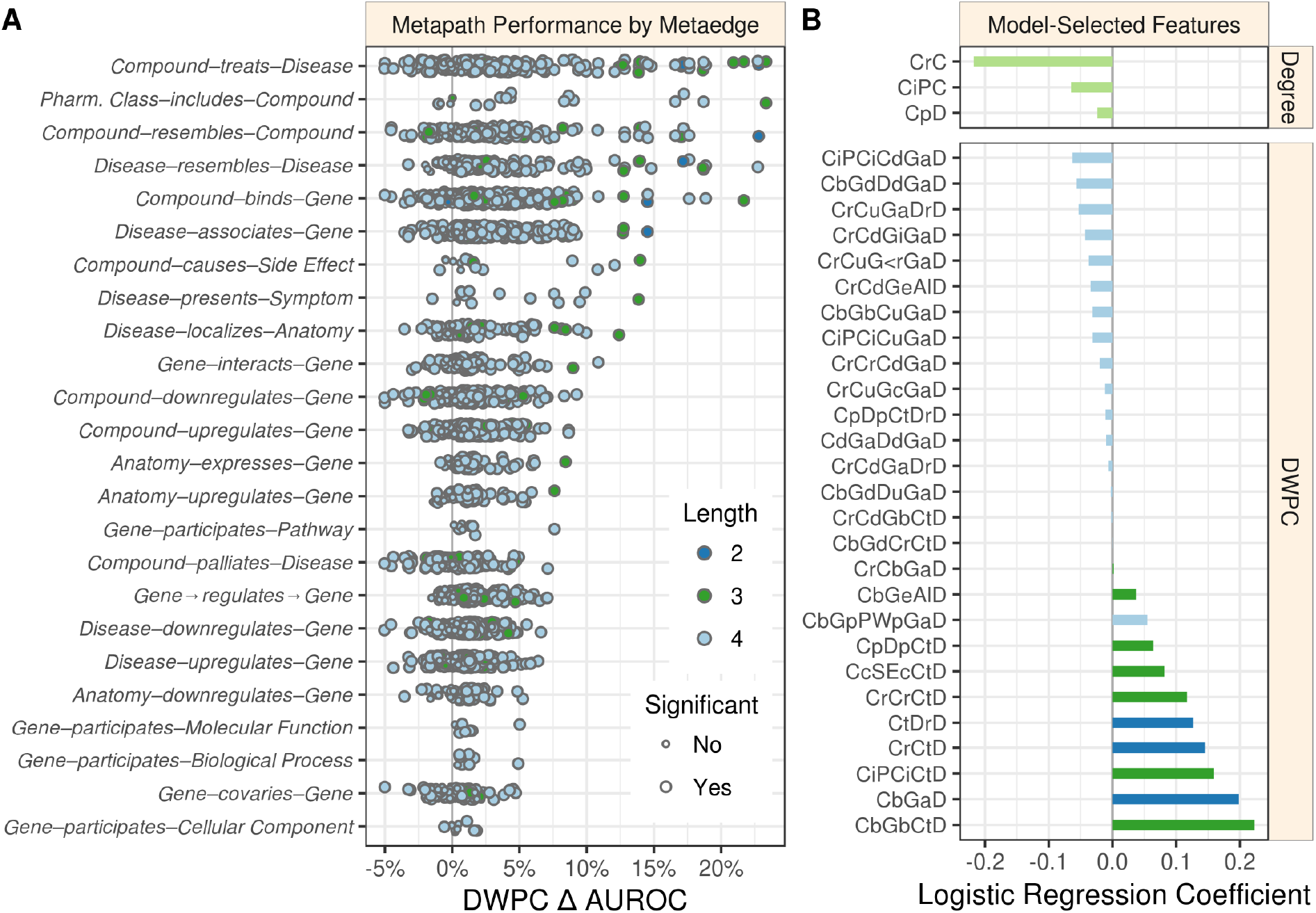
Performance by type and model coefficients. A) The performance of the DWPCs for 1,206 metapaths, organized by their composing metaedges. The larger dots represent metapaths that were significantly affected by permutation (false discovery rate < 5%). Metaedges are ordered by their best performing metapath. Since a metapath’s performance is limited by its least informative metaedge, the best performing metapath for a metaedge provides a lower bound on the pharmacologic utility of a given domain of information. B) Barplot of the model coefficients. Features were standardized prior to model fitting to make the coefficients comparable [39].

Overall, 709 of the 1,206 metapaths exhibited a statistically significant Δ AUROC at a false discovery rate cutoff of 5%. These 709 metapaths included all 24 metaedges, suggesting that each type of relationship we integrated provided at least some therapeutic utility. However, not all metaedges were equally present in significant metapaths: 259 significant metapaths included a *Compound–binds–Gene* metaedge, whereas only 4 included a *Gene–participates–Cellular Component* metaedge. Table 3 lists the predictiveness of several metapaths of interest. Refer to the Discussion for our interpretation of these findings.

**Table 3:** The predictiveness of select metapaths. A small selection of interesting or influential metapaths is provided (complete table online). Len. refers to number of metaedges composing the metapath. Δ AUROC and -log10(*p*) assess the performance of a metapath’s DWPC in discriminating treatments from non-treatments (in the all-features stage as described in Methods). *p* assesses whether permutation affected AUROC. For reference, *p* = 0.05 corresponds to -log10(*p*) = 1.30. Note that several metapaths shown here provided little evidence that Δ AUROC ≠ 0 underscoring their poor ability to predict whether a compound treated a disease. Coef. reports a metapath’s logistic regression coefficient as seen in Figure 2B. Metapaths removed in feature selection have missing coefficients whereas metapaths given zero-weight by the elastic net have coef. = 0.0.

| Abbrev. | Len. | $\Delta$<br>AUROC | $-\log_{10}(p)$ | Coef. | Metapath |
| --- | --- | --- | --- | --- | --- |
| CbGaD | 2 | 14.5% | 6.2 | 0.20 | Compound–binds–Gene–associates–Disease |
| CdGuD | 2 | 1.7% | 4.5 |  | Compound–downregulates–Gene–upregulates–Disease |
| CrCtD | 2 | 22.8% | 6.9 | 0.15 | Compound–resembles–Compound–treats–Disease |
| CtDrD | 2 | 17.2% | 5.8 | 0.13 | Compound–treats–Disease–resembles–Disease |
| CuGdD | 2 | 1.1% | 2.6 |  | Compound–upregulates–Gene–downregulates–Disease |
| CbGbCtD | 3 | 21.7% | 6.5 | 0.22 | Compound–binds–Gene–binds–Compound–treats–Disease |
| CbGeAlD | 3 | 8.4% | 5.2 | 0.04 | Compound–binds–Gene–expresses–Anatomy–localizes–Disease |
| CbGiGaD | 3 | 9.0% | 4.4 | 0.00 | Compound–binds–Gene–interacts–Gene–associates–Disease |
| CcSEcCtD | 3 | 14.0% | 6.8 | 0.08 | Compound–causes–Side Effect–causes–Compound–treats–Disease |
| CdGdCtD | 3 | 3.8% | 4.6 | 0.00 | Compound–downregulates–Gene–downregulates–Compound–treats–Disease |
| CdGuCtD | 3 | -2.1% | 2.4 |  | Compound–downregulates–Gene–upregulates–Compound–treats–Disease |
| CiPCiCtD | 3 | 23.3% | 7.5 | 0.16 | Compound–includes–Pharmacologic Class–includes–Compound–treats–Disease |
| CpDpCtD | 3 | 4.3% | 3.9 | 0.06 | Compound–palliates–Disease–palliates–Compound–treats–Disease |
| CrCrCtD | 3 | 17.0% | 5.0 | 0.12 | Compound–resembles–Compound–resembles–Compound–treats–Disease |
| CrCbGaD | 3 | 8.2% | 6.1 | 0.002 | Compound–resembles–Compound–binds–Gene–associates–Disease |
| CtDdGdD | 3 | 4.2% | 3.9 |  | Compound–treats–Disease–downregulates–Gene–downregulates–Disease |
| CtDdGuD | 3 | 0.5% | 1.0 |  | Compound–treats–Disease–downregulates–Gene–upregulates–Disease |
| CtDIAID | 3 | 12.4% | 6.0 |  | Compound–treats–Disease–localizes–Anatomy–localizes–Disease |
| CtDpSpD | 3 | 13.9% | 6.1 |  | Compound–treats–Disease–presents–Symptom–presents–Disease |
| CtDuGdD | 3 | 0.7% | 1.3 |  | Compound–treats–Disease–upregulates–Gene–downregulates–Disease |
| CtDuGuD | 3 | 1.1% | 1.4 |  | Compound–treats–Disease–upregulates–Gene–upregulates–Disease |
| CuGdCtD | 3 | -1.6% | 2.9 |  | Compound–upregulates–Gene–downregulates–Compound–treats–Disease |
| CuGuCtD | 3 | 4.4% | 3.5 | 0.00 | Compound–upregulates–Gene–upregulates–Compound–treats–Disease |
| CbGiGiGaD | 4 | 7.0% | 5.1 | 0.00 | Compound–binds–Gene–interacts–Gene–interacts–Gene–associates–Disease |
| CbGpBPpGaD | 4 | 4.9% | 3.8 | 0.00 | Compound–binds–Gene–participates–Biological Process–participates–Gene–associates–Disease |
| CbGpPWpGaD | 4 | 7.6% | 7.9 | 0.05 | Compound–binds–Gene–participates–Pathway–participates–Gene–associates–Disease |

### Predictions of drug efficacy

We implemented a machine learning approach to translate the network connectivity between a compound and a disease into a probability of treatment [40,41]. The approach relies on the 755 known treatments as positives and 29,044 non-treatments as negatives to train a logistic regression model. Note that 179,369 non-treatments were omitted as negative training observations because they had a prior probability of treatment equal to zero (see Methods). The features consisted of a prior probability of treatment, node degrees for 14 metaedges, and DWPCs for 123 metapaths that were well suited for modeling. A cross-validated elastic net was used to minimize overfitting, yielding a model with 31 features (Figure 2B). The DWPC features with negative coefficients appear to be included as node-degree-capturing covariates, i.e. they reflect the general connectivity of the compound and disease rather than specific paths between them. However, the 11 DWPC features with non-negligible positive coefficients represent the most salient types of connectivity for systematically modeling drug efficacy. See the metapaths with positive coefficients in Table 3 for unabbreviated names. As an example, the *CcSEcCtD* feature assesses whether the compound causes the same side effects as compounds that treat the disease. Alternatively, the *CbGeAlD* feature assesses whether the compound binds to genes that are expressed in the anatomies affected by the disease.

We applied this model to predict the probability of treatment between each of 1,538 connected compounds and each of 136 connected diseases, resulting in predictions for 209,168 compound– disease pairs [42], available at http://het.io/repurpose/. The 755 known disease-modifying indications were highly ranked (AUROC = 97.4%, Figure 3). The predictions also successfully prioritized two external validation sets: novel indications from DrugCentral (AUROC = 85.5%) and novel indications in clinical trial (AUROC = 70.0%). Together, these findings indicate that Project Rephetio has the ability to recognize efficacious compound–disease pairs.

**Figure 3:**
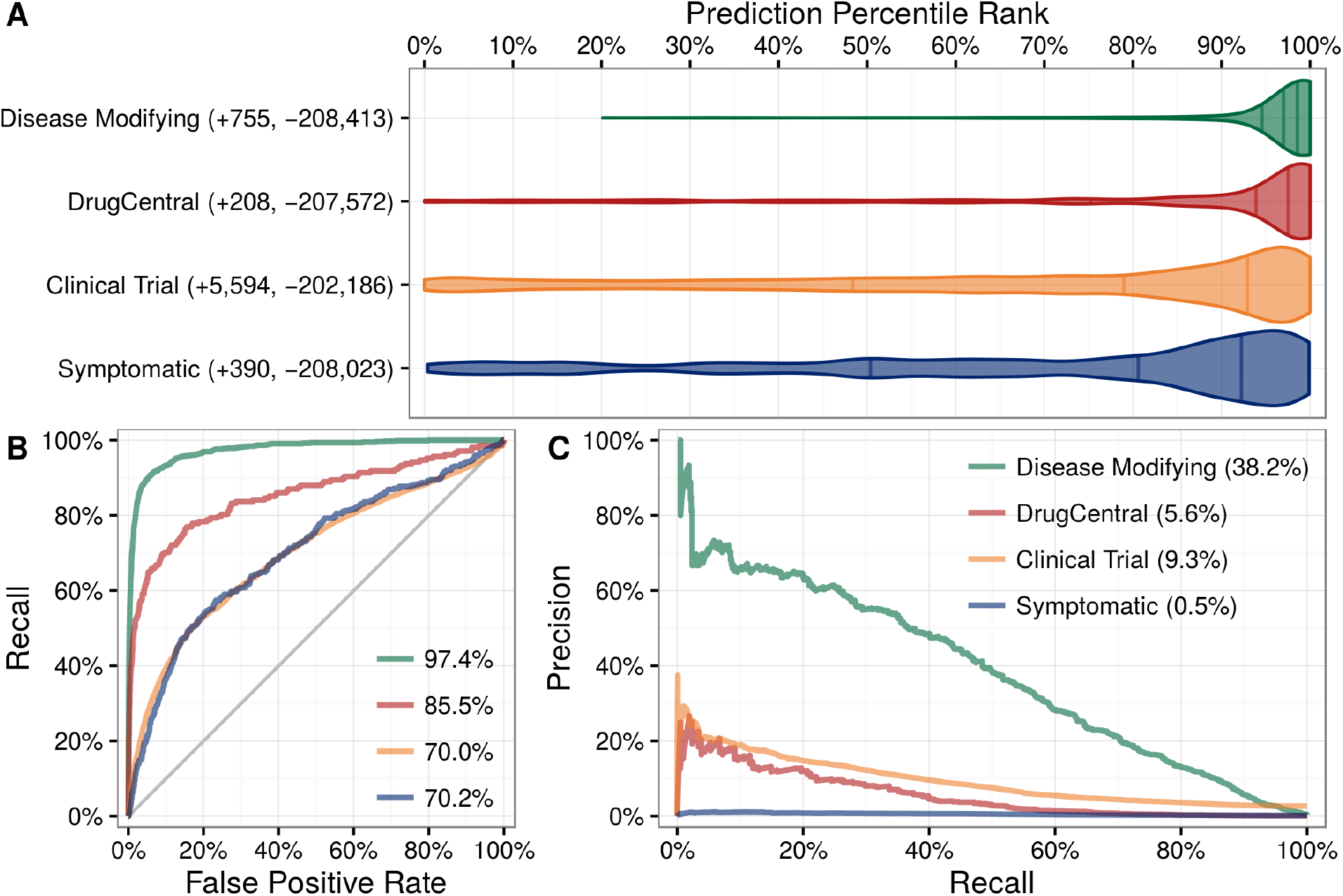
Predictions performance on four indication sets. We assess how well our predictions prioritize four sets of indications. A) The y-axis labels denote the number of indications (+) and non-indications (-) composing each set. Violin plots with quartile lines show the distribution of indications when compound–disease pairs are ordered by their prediction. In all four cases, the actual indications were ranked highly by our predictions. B) ROC Curves with AUROCs in the legend. C) Precision–Recall Curves with AUPRCs in the legend.

Predictions were scaled to the overall prevalence of treatments (0.36%). Hence a compound– disease pair that received a prediction of 1% represents a 2-fold enrichment over the null probability. Of the 3,980 predictions with a probability exceeding 1%, 586 corresponded to known disease-modifying indications, leaving 3,394 repurposing candidates. For a given compound or disease, we provide the percentile rank of each prediction. Therefore, users can assess whether a given prediction is a top prediction for the compound or disease. In addition, our table-based prediction browser links to a custom guide for each prediction, which displays in the Neo4j Hetionet Browser. Each guide includes a query to display the top paths supporting the prediction and lists clinical trials investigating the indication.

### Nicotine dependence case study

There are currently two FDA-approved medications for smoking cessation (varenicline and bupropion) that are not nicotine replacement therapies. PharmacotherapyDB v1.0 lists varenicline as a disease-modifying indication and nicotine itself as a symptomatic indication for nicotine dependence, but is missing bupropion. Bupropion was first approved for depression in 1985. Owing to the serendipitous observation that it decreased smoking in depressed patients taking this drug, Bupropion was approved for smoking cessation in 1997 [43]. Therefore we looked whether Project Rephetio could have predicted this repurposing. Bupropion was the 9th best prediction for nicotine dependence (99.5th percentile) with a probability 2.50-fold greater than the null. Figure 4 shows the top paths supporting the repurposing of bupropion.

**Figure 4:**
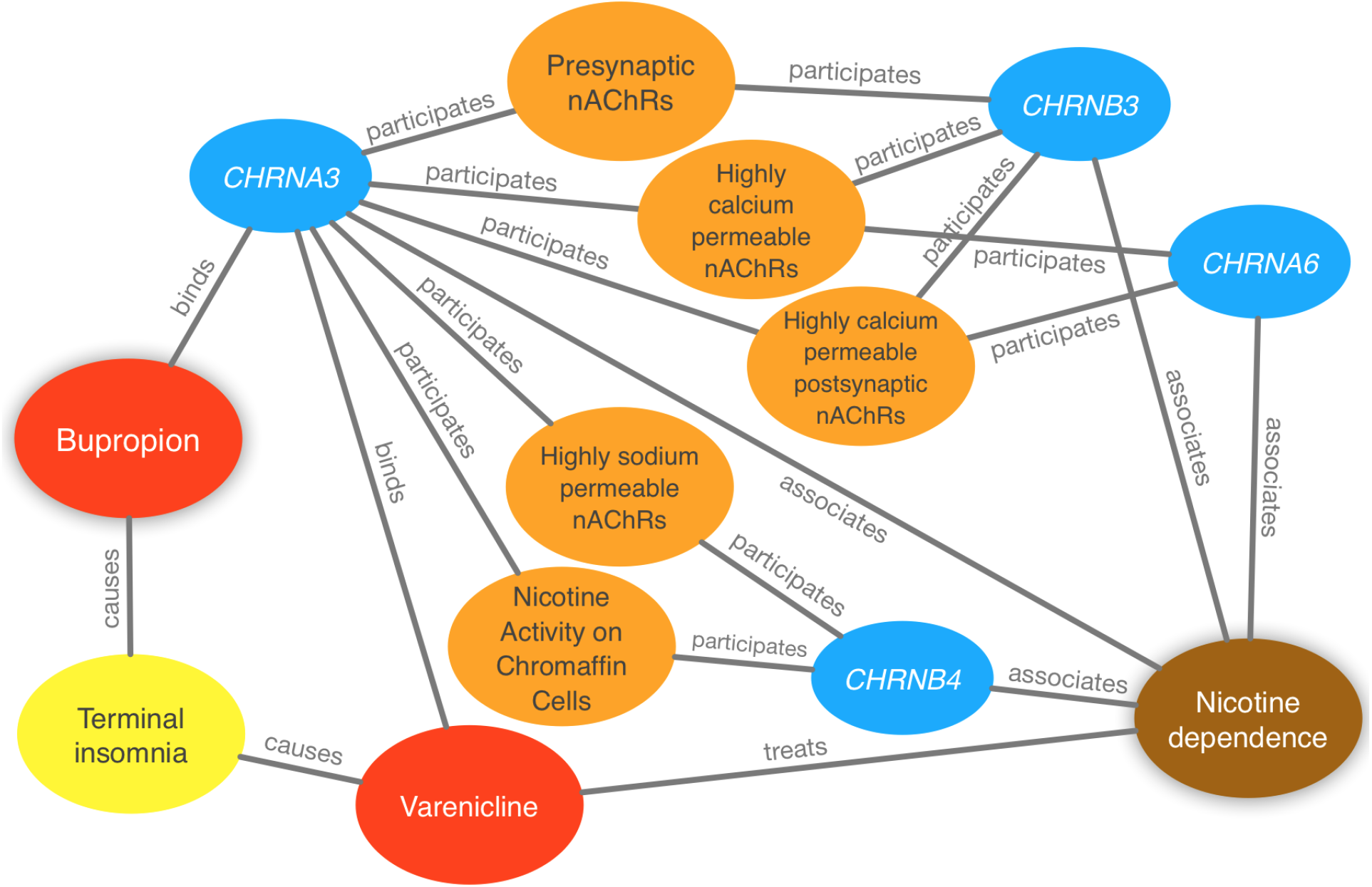
Evidence supporting the repurposing of bupropion for smoking cessation. This figure shows the 10 most supportive paths (out of 365 total) for treating nicotine dependence with bupropion, as available in this prediction’s Neo4j Browser guide. Our method detected that bupropion targets the CHRNA3 gene, which is also targeted by the known-treatment varenicline [44]. Furthermore, CHRNA3 is associated with nicotine dependence [45] and participates in several pathways that contain other nicotinic-acetylcholine-receptor (nAChR) genes associated with nicotine dependence. Finally, bupropion causes terminal insomnia [46] as does varenicline [47], which could indicate an underlying common mechanism of action.

Atop the nicotine dependence predictions were nicotine (10.97-fold over null), cytisine (10.58-fold), and galantamine (9.50-fold). Cytisine is widely used in Eastern Europe for smoking cessation due to its availability at a fraction of the cost of other pharmaceutical options [48]. In the last half decade, large scale clinical trials have confirmed cytisine’s efficacy [49,50]. Galantamine, an approved Alzheimer’s treatment, is currently in Phase 2 trial for smoking cessation and is showing promising results [51]. In summary, nicotine dependence illustrates Project Rephetio’s ability to predict efficacious treatments and prioritize historic and contemporary repurposing opportunities.

### Epilepsy case study

Several factors make epilepsy an interesting disease for evaluating repurposing predictions [52]. Antiepileptic drugs work by increasing the seizure threshold — the amount of electric stimulation that is required to induce seizure. The effect of a drug on the seizure threshold can be cheaply and reliably tested in rodent models. As a result, the viability of most approved drugs in treating epilepsy is known.

We focused our evaluation on the top 100 scoring compounds — referred to as the epilepsy predictions in this section — after discarding a single combination drug. We classified each compound as anti-ictogenic (seizure suppressing), unknown (no established effect on the seizure threshold), or ictogenic (seizure generating) according to medical literature [52]. Of the top 100 epilepsy predictions, 77 were anti-ictogenic, 8 were unknown, and 15 were ictogenic (Figure 5A). Notably, the predictions contained 23 of the 25 disease-modifying antiepileptics in PharamcotherapyDB v1.0.

**Figure 5:**
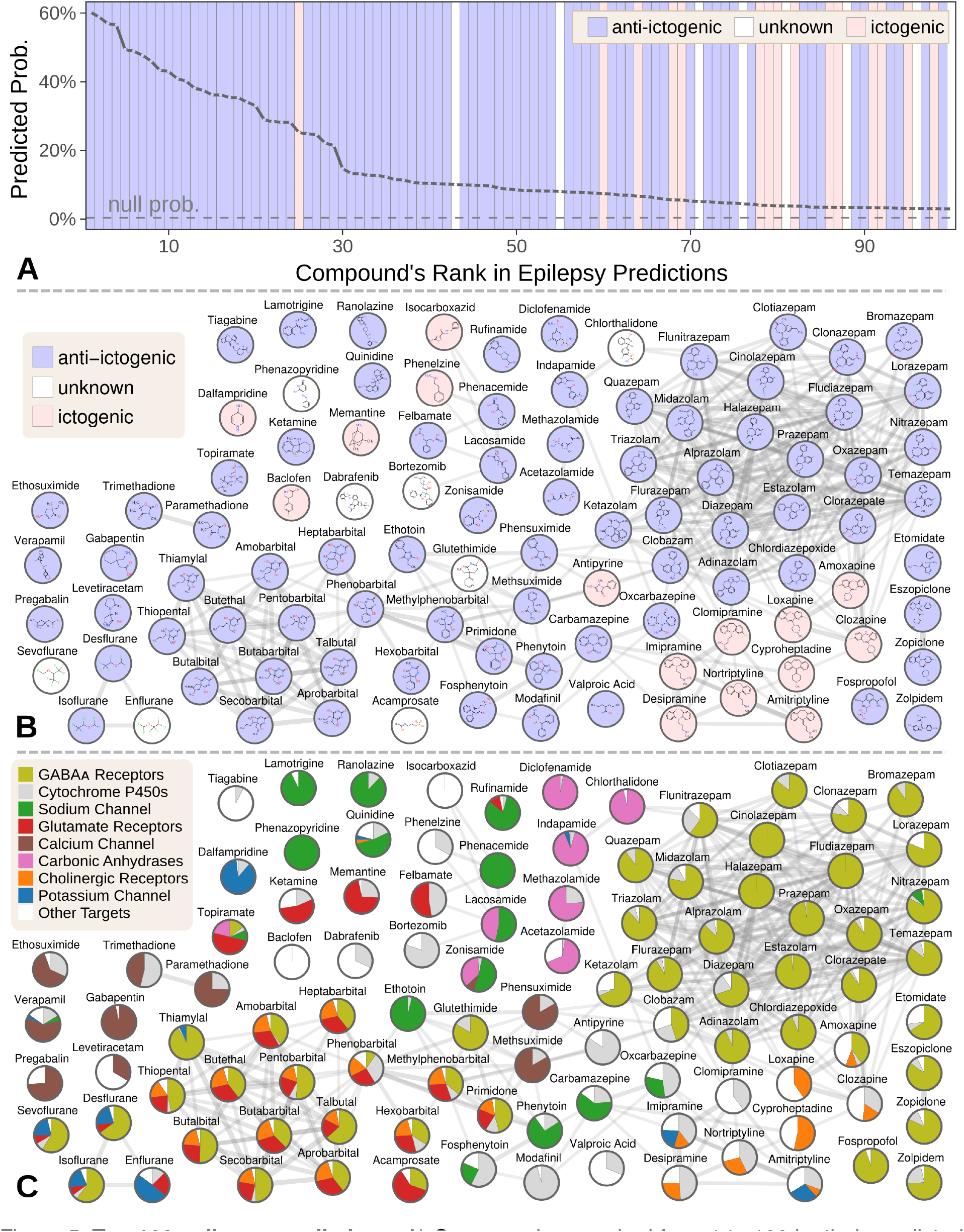
Top 100 epilepsy predictions. A) Compounds — ranked from 1 to 100 by their predicted probability of treating epilepsy — are colored by their effect on seizures [52]. The highest predictions are almost exclusively anti-ictogenic. Further down the prediction list, the prevalence of drugs with an ictogenic (contraindication) or unknown (novel repurposing candidate) effect on epilepsy increases. All compounds shown received probabilities far exceeding the null probability of treatment (0.36%). B) A chemical similarity network of the epilepsy predictions, with each compound’s 2D structure [53]. Edges are *Compound–resembles–Compound* relationships from Hetionet v1.0. Nodes are colored by their effect on seizures. C) The relative contribution of important drug targets to each epilepsy prediction [53]. Specifically, pie charts show how the 8 most-supportive drug targets across all 100 epilepsy predictions contribute to individual predictions. Other Targets represents the aggregate contribution of all targets not listed. The network layout is identical to B.

Many of the 77 anti-ictogenic compounds were not first-line antiepileptic drugs. Instead, they were used as ancillary drugs in the treatment of status epilepticus. For example, we predicted four halogenated ethers, two of which (isoflurane and desflurane) are used clinically to treat life-threatening seizures that persist despite treatment [54]. As inhaled anesthetics, these compounds are not appropriate as daily epilepsy medications, but are feasible for refractory status epilepticus where patients are intubated.

Given this high precision (77%), the 8 compounds of unknown effect are promising repurposing candidates. For example, acamprosate — whose top prediction was epilepsy — is a taurine analog that promotes alcohol abstinence. Support for this repurposing arose from acamprosate’s inhibition of the glutamate receptor and positive modulation of the GABAᴀ receptor (Figure 5C). If effective against epilepsy, acamprosate could serve a dual benefit for recovering alcoholics who experience seizures from alcohol withdrawal.

While certain classes of compounds were highly represented in our epilepsy predictions, such benzodiazepines and barbiturates, there was also considerable diversity [52]. The 100 predicted compounds encompassed 26 third-level ATC codes [55], such as antiarrhythmics (quinidine, classified as anti-ictogenic) and urologicals (phenazopyridine, classified as unknown). Furthermore, 25 of the compounds were chemically distinct, i.e. they did not resemble any of the other epilepsy predictions (Figure 5B).

Next, we investigated which components of Hetionet contributed to the epilepsy predictions [52]. In total, 392,956 paths of 12 types supported the predictions. Using several different methods for grouping paths, we were able to quantify the aggregate biological evidence. Our algorithm primarily drew on two aspects of epilepsy: its known treatments (76% of the total support) and its genetic associations (22% of support). In contrast, our algorithm drew heavily on several aspects of the predicted compounds: their targeted genes (44%), their chemically similar compounds (30%), their pharmacologic classes, their palliative indications (5%), and their side effects (4%).

Specifically, 266,192 supporting paths originated with a Compound–binds–Gene relationship. Aggregating support by these genes shows the extent that 121 different drug targets contributed to the predictions [52]. In order of importance, the predictions targeted GABAᴀ receptors (15.3% of total support), cytochrome P450 enzymes (5.6%), the sodium channel (4.6%), glutamate receptors (3.8%), the calcium channel (2.7%), carbonic anhydrases (2.5%), cholinergic receptors (2.1%) and the potassium channel (1.4%). Besides cytochrome P450, which primarily influences pharmacokinetics [56], our method detected and leveraged bonafide anti-ictogenic mechanisms [57]. Figure 5C shows drug target contributions per compound and illustrates the considerable mechanistic diversity among the predictions.

Also notable are the 15 ictogenic compounds in our top 100 predictions. Nine of the ictogenic compounds share a tricyclic structure (Figure 5B), five of which are tricyclic antidepressants. While the ictogenic mechanisms of these antidepressants are still unclear [58], Figure 5C suggests their anticholinergic effects may be responsible [59], in accordance with previous theories [60].

We also ranked the contribution of the 1,137 side effects that supported the epilepsy predictions through 117,720 CcSEcCtD paths. The top five side effects — ataxia (0.069% of total support), nystagmus (0.049%), diplopia (0.045%), somnolence (0.044%), and vomiting (0.043%) — reflect established adverse effects of antiepileptic drugs [61–65]. In summary, our method simultaneously identified the hallmark side effects of antiepileptic drugs while incorporating this knowledge to prioritize 1,538 compounds for anti-ictogenic activity.

## Discussion

We created Hetionet v1.0 by integrating 29 resources into a single data structure — the hetnet. Consisting of 11 types of nodes and 24 types of relationships, Hetionet v1.0 brings more types of information together than previous leading-studies in biological data integration [66]. Moreover, we strove to create a reusable, extensible, and property-rich network. While all of the resources we include are publicly available, their integration was a time-intensive undertaking. Hetionet allows researchers to begin answering integrative questions without having to first spend months processing data.

Our public Neo4j instance allows users to immediately interact with Hetionet. Through the Cypher language, users can perform highly specialized graph queries with only a few lines of code. Queries can be executed in the web browser or programmatically from a language with a Neo4j driver. For users that are unfamiliar with Cypher, we include several example queries in a Browser guide. In contrast to traditional REST APIs, our public Neo4j instance provides users with maximal flexibility to construct custom queries by exposing the underlying database.

As data has grown more plentiful and diverse, so has the applicability of hetnets. Unfortunately, network science has been naturally fragmented by discipline resulting in relatively slow progress in integrating heterogeneous data. A 2014 analysis identified 78 studies using multilayer networks — a superset of hetnets (heterogeneous information networks) with the potential for additional dimensions, such as time. However, the studies relied on 26 different terms, 9 of which had multiple definitions [67,68]. Nonetheless, core infrastructure and algorithms for hetnets are emerging. Compared to the existing mathematical frameworks for multilayer networks that must deal with layers other than type (such as the aspect of time) [67], the primary obligation of hetnet algorithms is to be type aware. One goal of our project has been to unite hetnet research across disciplines. We approached this goal by making Project Rephetio entirely available online and inviting community feedback throughout the process [69].

Integrating every resource into a single interconnected data structure allowed us to assess systematic mechanisms of drug efficacy. Using the max performing metapath to assess the pharmacological utility of a metaedge (Figure 2A), we can divide our relationships into tiers of informativeness. The top tier consists of the types of information traditionally considered by pharmacology: *Compound–treats–Disease*, *Pharmacologic Class–includes–Compound*, *Compound–resembles–Compound*, *Disease–resembles–Disease*, and *Compound–binds–Gene*. The upper-middle tier consists of types of information that have been the focus of substantial medical study, but have only recently started to play a bigger role in drug development, namely the metaedges *Disease–associates–Gene*, *Compound–causes–Side Effect*, *Disease–presents– Symptom*, *Disease–localizes–Anatomy*, and *Gene–interacts–Gene*.

The lower-middle tier contains the transcriptomics metaedges such as *Compound–downregulates– Gene*, *Anatomy–expresses–Gene*, *Gene→regulates→Gene*, and *Disease–downregulates–Gene*. Much excitement surrounds these resources due to their high throughput and genome-wide scope, which offers a route to drug discovery that is less biased by existing knowledge. However, our findings suggest that these resources are only moderately informative of drug efficacy. Other lower-middle tier metaedges were the product of time-intensive biological experimentation, such as *Gene–participates–Pathway*, *Gene–participates–Molecular Function*, and *Gene–participates– Biological Process*. Unlike the top tier resources, this knowledge has historically been pursued for basic science rather than primarily medical applications. The weak yet appreciable performance of the *Gene–covaries–Gene* suggests the synergy between the fields of evolutionary genomics and disease biology. The lower tier included the *Gene–participates–Cellular Component* metaedge, which may reflect that the relevance of cellular location to pharmacology is highly case dependent and not amenable to systematic profiling.

The performance of specific metapaths (Table 3) provides further insight. For example, significant emphasis has been put on the use of transcriptional data for drug repurposing [30]. One common approach has been to identify compounds with opposing transcriptional signatures to a disease [18,70]. However, several systematic studies report underwhelming performance of this approach [24–26] — a finding supported by the low performance of the *CuGdD* and *CdGuD* metapaths in Project Rephetio. Nonetheless, other transcription-based methods showed some promise. Compounds with similar transcriptional signatures were prone to treating the same disease (*CuGuCtD* and *CdGdCtD* metapaths), while compounds with opposing transcriptional signatures were slightly averse to treating the same disease (*CuGdCtD* and *CdGuCtD* metapaths). In contrast, diseases with similar transcriptional profiles were not prone to treatment by the same compound (*CtDdGuD* and *CtDuGdD*).

By comparably assessing the informativeness of different metaedges and metapaths, Project Rephetio aims to guide future research towards promising data types and analyses. One caveat is that omics-scale experimental data will likely play a larger role in developing the next generation of pharmacotherapies. Hence, were performance reevaluated on treatments discovered in the forthcoming decades, the predictive ability of these data types may rise. Encouragingly, most data types were at least weakly informative and hence suitable for further study. Ideally, different data types would provide orthogonal information. However, our model for whether a compound treats a disease focused on 11 metapaths — a small portion of the hundreds of metapaths available. While parsimony aids interpretation, our model did not draw on the weakly-predictive high-throughput data types — which are intriguing for their novelty, scalability, and cost-effectiveness — as much as we had hypothesized.

Instead our model selected types of information traditionally considered in pharmacology. However unlike a pharmacologist whose area of expertise may be limited to a few drug classes, our model was able to predict probabilities of treatment for all 209,168 compound–disease pairs. Furthermore, our model systematically learned the importance of each type of network connectivity. For any compound–disease pair, we now can immediately provide the top network paths supporting its therapeutic efficacy. A traditional pharmacologist may be able to produce a similar explanation, but likely not until spending substantial time researching the compound’s pharmacology, the disease’s pathophysiology, and the molecular relationships in between. Accordingly, we hope certain predictions will spur further research, such as trials to investigate the off-label use of acamprosate for epilepsy, which is supported by one animal model [71].

As demonstrated by the 15 ictogenic compounds in our top 100 epilepsy predictions, Project Rephetio’s predictions can include contraindications in addition to indications. Since many of Hetionet v1.0’s relationship types are general (e.g. the *Compound–binds–Gene* relationship type conflates antagonist with agonist effects), we expect some high scoring predictions to exacerbate rather than treat the disease. However, the predictions made by Hetionet v1.0 represent such substantial relative enrichment over the null that uncovering the correct directionality is a logical next step and worth undertaking. Going forward, advances in automated mining of the scientific literature could enable extraction of precise relationship types at omics scale [72,73].

Future research should focus on gleaning orthogonal information from data types that are so expansive that computational methods are the only option. Our *CuGuCtD* feature — measuring whether a compound upregulates the same genes as compounds which treat the disease — is a good example. This metapath was informative by itself (Δ AUROC = 4.4%) but was not selected by the model, despite its orthogonal origin (gene expression) to selected metapaths. Using a more extensive catalog of treatments as the gold standard would be one possible approach to increase the power of feature selection.

Integrating more types of information into Hetionet should also be a future priority. The “network effect” phenomenon suggests that the addition of each new piece of information will enhance the value of Hetionet’s existing information. We envision a future where all biological knowledge is encoded into a single hetnet. Hetionet v1.0 was an early attempt, and we hope the strong performance of Project Rephetio in repurposing drugs foreshadows the many applications that will thrive from encoding biology in hetnets.

## Methods

Hetionet was built entirely from publicly available resources with the goal of integrating a broad diversity of information types of medical relevance, ranging in scale from molecular to organismal. Practical considerations such as data availability, licensing, reusability, documentation, throughput, and standardization informed our choice of resources. We abided by a simple litmus test for determining how to encode information in a hetnet: nodes represent nouns, relationships represent verbs [74,75].

Our method for relationship prediction creates a strong incentive to avoid redundancy, which increases the computational burden without improving performance. In a previous study to predict disease–gene associations using a hetnet of pathophysiology [22], we found that different types of gene sets contributed highly redundant information. Therefore, in Hetionet v1.0 we reduced the number of gene set node types from 14 to 3 by omitting several gene set collections and aggregating all pathway nodes.

### Nodes

Nodes encode entities. We extracted nodes from standard terminologies, which provide curated vocabularies to enable data integration and prevent concept duplication. The ease of mapping external vocabularies, adoption, and comprehensiveness were primary selection criteria. Hetionet v1.0 includes nodes from 5 ontologies — which provide hierarchy of entities for a specific domain — selected for their conformity to current best practices [76].

We selected 137 terms from the Disease Ontology [77,78] (which we refer to as DO Slim [79,80]) as our **disease** set. Our goal was to identify complex diseases that are distinct and specific enough to be clinically relevant yet general enough to be well annotated. To this end, we included diseases that have been studied by GWAS and cancer types from `TopNodes_DOcancerslim` [81]. We ensured that no DO Slim disease was a subtype of another DO Slim disease. **Symptoms** were extracted from MeSH by taking the 438 descendants of *Signs and Symptoms* [82,83].

Approved small molecule **compounds** with documented chemical structures were extracted from DrugBank version 4.2 [84–86]. Unapproved compounds were excluded because our focus was repurposing. In addition, unapproved compounds tend to be less studied than approved compounds making them less attractive for our approach where robust network connectivity is critical. Finally, restricting to small molecules with known documented structures enabled us to map between compound vocabularies (see Mappings).

**Side effects** were extracted from SIDER version 4.1 [87–89]. SIDER codes side effects using UMLS identifiers [90], which we also adopted. **Pharmacologic Classes** were extracted from the DrugCentral data repository [91,. Only pharmacologic classes corresponding to the “Chemical/Ingredient”, “Mechanism of Action”, and “Physiologic Effect” FDA class types were included to avoid pharmacologic classes that were synonymous with indications [92].

Protein-coding human **genes** were extracted from Entrez Gene [93–95]. Anatomical structures, which we refer to as **anatomies**, were extracted from Uberon [96]. We selected a subset of 402 Uberon terms by excluding terms known not to exist in humans and terms that were overly broad or arcane [97,98].

**Pathways** were extracted by combining human pathways from WikiPathways [99,100], Reactome [101], and the Pathway Interaction Database [102]. The latter two resources were retrieved from Pathway Commons [103], which compiles pathways from several providers. Duplicate pathways and pathways without multiple participating genes were removed [104,105]. **Biological processes, cellular components**, and **molecular functions** were extracted from the Gene Ontology [106]. Only terms with 2–1000 annotated genes were included.

### Mappings

Before adding relationships, all identifiers needed to be converted into the vocabularies matching that of our nodes. Oftentimes, our node vocabularies included external mappings. For example, the Disease Ontology includes mappings to MeSH, UMLS, and the ICD, several of which we submitted during the course of this study [107]. In a few cases, the only option was to map using gene symbols, a disfavored method given that it can lead to ambiguities.

When mapping external disease concepts onto DO Slim, we used transitive closure. For example, the UMLS concept for primary progressive multiple sclerosis (`C0751964`) was mapped to the DO Slim term for multiple sclerosis (`DOID:2377`).

Chemical vocabularies presented the greatest mapping challenge [85], since these are poorly standardized [108]. UniChem’s [109] Connectivity Search [110] was used to map compounds, which maps by atomic connectivity (based on First InChIKey Hash Blocks [111]) and ignores small molecular differences.

### Edges

*Anatomy–downregulates–Gene* and *Anatomy–upregulates–Gene* edges [112–114] were extracted from Bgee [115], which computes differentially expressed genes by anatomy in post-juvenile adult humans. *Anatomy–expresses–Gene* edges were extracted from Bgee and TISSUES [116–118].

Compound–binds–Gene edges were aggregated from BindingDB [119,120], DrugBank [84,121], and DrugCentral [91]. Only binding relationships to single proteins with affinities of at least 1 μM (as determined by Kd, K_i_, or IC_50_) were selected from the October 2015 release of BindingDB [122, 123]. Target, carrier, transporter, and enzyme interactions with single proteins (i.e. excluding protein groups) were extracted from DrugBank 4.2 [86,124]. In addition, all mapping DrugCentral target relationships were included [92].

*Compound–treats–Disease* (disease-modifying indications) and *Compound–palliates–Disease* (symptomatic indications) edges are from PharmacotherapyDB as described in Intermediate resources. *Compound–causes–Side Effect* edges were obtained from SIDER 4.1 [87–89], which uses natural language processing to identify side effects in drug labels. *Compound–resembles– Compound* relationships [86,126] represent chemical similarity and correspond to a Dice coefficient ≥ 0.5 [127] between extended connectivity fingerprints [128,129]. *Pharmacologic Class– includes–Compound* edges were extracted from DrugCentral for three FDA class types [91, 92]. *Compound–downregulates–Gene* and *Compound–upregulates–Gene* relationships were computed from LINCS L1000 as described in Intermediate resources.

*Disease–associates–Gene edges* were extracted from the GWAS Catalog [130], DISEASES [131, 132], DisGeNET [133,134], and DOAF [135,136]. The GWAS Catalog compiles disease–SNP associations from published GWAS [137]. We aggregated overlapping loci associated with each disease and identified the mode reported gene for each high confidence locus [138,139]. DISEASES integrates evidence of association from text mining, curated catalogs, and experimental data [140]. Associations from DISEASES with integrated scores ≥ 2 were included after removing the contribution of DistiLD. DisGeNET integrates evidence from over 10 sources and reports a single score for each association [141,142]. Associations with scores ≥ 0.06 were included. DOAF mines Entrez Gene GeneRIFs (textual annotations of gene function) for disease mentions [143]. Associations with 3 or more supporting GeneRIFs were included. *Disease–downregulates–Gene and Disease–upregulates–Gene* relationships [144,145] were computed using STARGEO as described in Intermediate resources.

*Disease–localizes–Anatomy, Disease–presents–Symptom*, and Disease–resembles–Disease edges were calculated from MEDLINE co-occurrence [82,146]. MEDLINE is a subset of 21 million PubMed articles for which designated human curators have assigned topics. When retrieving articles for a given topic (MeSH term), we activated two non-default search options as specified below: `majr` for selecting only articles where the topic is major and `noexp` for suppressing explosion (returning articles linked to MeSH subterms). We identified 4,161,769 articles with two or more disease topics; 696,252 articles with both a disease topic (`majr`) and an anatomy topic (`noexp`) [147]; and 363,928 articles with both a disease topic (`majr`) and a symptom topic (`noexp`). We used a Fisher’s exact test [148] to identify pairs of terms that occurred together more than would be expected by chance in their respective corpus. We included co-occurring terms with p < 0.005 in Hetionet v1.0.

*Gene→regulates→Gene* directed edges were generated from the LINCS L1000 genetic interference screens (see Intermediate resources) and indicate that knockdown or overexpression of the source gene significantly dysregulated the target gene [149,150]. Gene–covaries–Gene edges represent evolutionary rate covariation ≥ 0.75 [151–153]. Gene–interacts–Gene edges [154, 135] represent when two genes produce physically-interacting proteins. We compiled these interactions from the Human Interactome Database [156–159], the Incomplete Interactome [160], and our previous study [22]. *Gene–participates–Biological Process, Gene–participates–Cellular Component*, and *Gene–participates–Molecular* Function edges are from Gene Ontology annotations [161]. As described in Intermediate resources, annotations were propagated [162,163]. *Gene–participates–Pathway edges* were included from the human pathway resources described in the Nodes section [104, 105].

### Directionality

Whether a certain type of relationship has directionality is defined at the metaedge level. Directed metaedges are only necessary when they connect a metanode to itself and correspond to an asymmetric relationship. In the case of Hetionet v1.0, the sole directed metaedge was *Gene→regulates→Gene*. To demonstrate the implications of directionality, Hetionet v1.0 contains two relationships between the genes *HADH* and *STAT1: HADH–interacts–STAT1* and *HADH→regulates→STAT1*. Both edges can be represented in the inverse orientation: *STAT1– interacts–HADH* and *STAT1←regulates←HADH*. However due to directed nature of the regulates relationship, STAT1→regulates→HADH is a distinct edge, which does not exist in the network. Similarly, *HADH–associates–obesity* and *obesity–associates–HADH* are inverse orientations of the same underlying undirected relationship. Accordingly, the following path exists in the network: *obesity–associates–HADH→regulates→STAT1*, which can also be inverted to *STAT1←regulates←HADH–associates–obesity*.

### Intermediate resources

In the process of creating Hetionet, we produced several datasets with broad applicability that extended beyond Project Rephetio. These resources are referred to as intermediate resources and described below.

### Transcriptional signatures of disease using STARGEO

STARGEO is a nascent platform for annotating and meta-analyzing differential gene expression experiments. The STAR acronym stands for Search-Tag-Analyze Resources, while GEO refers to the Gene Expression Omnibus [164,165]. STARGEO is a layer on top of GEO that crowdsources sample annotation and automates meta-analysis.

Using STARGEO, we computed differentially expressed genes between healthy and diseased samples for 49 diseases [144,145]. First, we and others created case/control tags for 66 diseases. After combing through GEO series and tagging samples, 49 diseases had sufficient data for case-control meta-analysis: multiple series with at least 3 cases and 3 controls. For each disease, we performed a random effects meta-analysis on each gene to combine log2 fold-change across series. These analyses incorporated 27,019 unique samples from 460 series on 107 platforms.

Differentially expressed genes (false discovery rate ≤ 0.05) were identified for each disease. The median number of upregulated genes per disease was 351 and the median number of downregulated genes was 340. Endogenous depression was the only of the 49 diseases without any significantly dysregulated genes.

### Transcriptional signatures of perturbation from LINCS L1000

LINCS L1000 profiled the transcriptional response to small molecule and genetic interference perturbations. To increase throughput, expression was only measured for 978 genes, which were selected for their ability to impute expression of the remaining genes. A single perturbation was often assayed under a variety of conditions including cell types, dosages, timepoints, and concentrations. Each condition generates a single signature of dysregulation z-scores. We further processed these signatures to fit into our approach [166,167].

First we computed consensus signatures — which meta-analyze multiple signatures to condense them into one — for DrugBank small molecules, Entrez genes, and all L1000 perturbations [149, 150]. First, we discarded non-gold (non-replicating or indistinct) signatures. Then we meta-analyzed z-scores using Stouffer’s method. Each signature was weighted by its average Spearman’s correlation to other signatures, with a 0.05 minimum, to de-emphasize discordant signatures. Our signatures include the 978 measured genes and the 6,489 imputed genes from the “best inferred gene subset”. To identify significantly dysregulated genes, we selected genes using a Bonferroni cutoff of p = 0.05 and limited the number of imputed genes to 1,000.

The consensus signatures for genetic perturbations allowed us to assess various characteristics of the L1000 dataset. First, we looked at whether genetic interference dysregulated its target gene in the expected direction [168]. Looking at measured z-scores for target genes, we found that the knockdown perturbations were highly reliable, while the overexpression perturbations were only moderately reliable with 36% of overexpression perturbations downregulating their target. However, imputed z-scores for target genes barely exceeded chance at responding in the expected direction to interference. Hence, we concluded that the imputation quality of LINCS L1000 is poor. However, when restricting to significantly dyseregulated targets, 22 out of 29 imputed genes responded in the expected direction. This provides some evidence that the directional fidelity of imputation is higher for significantly dysregulated genes. Finally, we found that the transcriptional signatures of knocking down and overexpressing the same gene were positively correlated 65% of the time, suggesting the presence of a general stress response [169].

Based on these findings, we performed additional filtering of signifcantly dysregulated genes when building Hetionet v1.0. *Compound–down/up-regulates–Gene* relationships were restricted to the 125 most significant per compound-direction-status combination (status refers to measured versus imputed). For genetic interference perturbations, we restricted to the 50 most significant genes per gene-direction-status combination and merged the remaining edges into a single *Gene→regulates→Gene* relationship type containing both knockdown and overexpression perturbations.

### PharmacotherapyDB: physician curated indications

We created PharmacotherapyDB, an open catalog of drug therapies for disease [170–172]. Version 1.0 contains 755 disease-modifying therapies and 390 symptomatic therapies between 97 diseases and 601 compounds.

This resource was motivated by the need for a gold standard of medical indications to train and evaluate our approach. Initially, we identified four existing indication catalogs [173]: MEDI-HPS which mined indications from RxNorm, SIDER 2, MedlinePlus, and Wikipedia [174]; LabeledIn which extracted indications from drug labels via human curation [175–177]; EHRLink which identified medication–problem pairs that clinicians linked together in electronic health records [178, 179]; and indications from PREDICT, which were compiled from UMLS relationships, drugs.com, and drug labels [24]. After mapping to DO Slim and DrugBank Slim, the four resources contained 1,388 distinct indications.

However, we noticed that many indications were palliative and hence problematic as a gold standard of pharmacotherapy for our in *silico* approach. Therefore, we recruited two practicing physicians to curate the 1,388 preliminary indications [180]. After a pilot on 50 indications, we defined three classifications: *disease modifying* meaning a drug that therapeutically changes the underlying or downstream biology of the disease; *symptomatic* meaning a drug that treats a significant symptom of the disease; and *non-indication* meaning a drug that neither therapeutically changes the underlying or downstream biology nor treats a significant symptom of the disease. Both curators independently classified all 1,388 indications.

The two curators disagreed on 444 calls (Cohen’s κ = 49.9%). We then recruited a third practicing physician, who reviewed all 1,388 calls and created a detailed explanation of his methodology [180]. We proceeded with the third curator’s calls as the consensus curation. The first two curators did have reservations with classifying steroids as disease modifying for autoimmune diseases. We ultimately considered that these indications met our definition of disease modifying, which is based on a pathophysiological rather than clinical standard. Accordingly, therapies we consider disease modifying may not be used to alter long-term disease course in the modern clinic due to a poor risk–benefit ratio.

### User-friendly Gene Ontology annotations

We created a browser (http://git.dhimmel.com/gene-ontology/) to provide straightforward access to Gene Ontology annotations [162,163]. Our service provides annotations between Gene Ontology terms and Entrez Genes. The user chooses propagated/direct annotation and all/experimental evidence. Annotations are currently available for 37 species and downloadable as user-friendly TSV files.

### Data copyright and licensing

We committed to openly releasing our data and analyses from the origin of the project [181]. Our goals were to contribute to the advancement of science [182,183], maximize our impact [184,185], and enable reproducibility [186–188]. These objectives required publicly distributing and openly licensing Hetionet and Project Rephetio data and analyses [189,190].

Since we integrated only public resources, which were overwhelmingly funded by academic grants, we had assumed that our project and open sharing of our network would not be an issue. However, upon releasing a preliminary version of Hetionet [191], a community reviewer informed us of legal barriers to integrating public data. In essence, both copyright (rights of exclusivity automatically granted to original works) and terms of use (rules that users must agree to in order to use a resource) place legally-binding restrictions on data reuse. In short, public data is not by default open data.

Hetionet v1.0 integrates 29 resources, but two resources were removed prior to the v1.0 release. Of the total 31 resources [192], five were United States government works not subject to copyright, and twelve had licenses that met the Open Definition of knowledge version 2.1. Four resources allowed only non-commercial reuse. Most problematic were the remaining nine resources that had no license — which equates to all rights reserved by default and forbids reuse [193] — and one resource that explicitly forbid redistribution.

Additional difficulty resulted from license incompatibles across resources, which was caused primarily by non-commercial and share-alike stipulations. Furthermore, it was often unclear who owned the data [194]. Therefore, we sought input from legal experts and chronicled our progress [192,–198].

Ultimately, we did not find an ideal solution. We had to choose between absolute compliance and Hetionet: strictly adhering to copyright and licensing arrangements would have decimated the network. On the other hand, in the United States, mere facts are not subject to copyright, and fair use doctrine helps protect reuse that is transformative and educational. Hence, we choose a path forward which balanced legal, normative, ethical, and scientific considerations.

If a resource was in the public domain, we licensed any derivatives as CC0 1.0. For resources licensed to allow reuse, redistribution, and modification, we transmitted their licenses as properties on the specific nodes and relationships in Hetionet v1.0. For all other resources — for example, resources without licenses or with licenses that forbid redistribution — we sent permission requests to their creators. The median time till first response to our permission requests was 16 days, with only 2 resources affirmatively granting us permission. We did not receive any responses asking us to remove a resource. However, we did voluntarily remove MSigDB [199], since its license was highly problematic [195]. As a result of our experience, we recommend that publicly-funded data should be explicitly dedicated to the public domain whenever possible.

### Permuted hetnets

From Hetionet, we derived five permuted hetnets [200]. The permutations preserve node degree but eliminate edge specificity by employing an algorithm called XSwap to randomly swap edges [201]. To extend XSwap to hetnets [22], we permuted each metaedge separately, so that edges were only swapped with other edges of the same type. We adopted a Markov chain approach, whereby the first permuted hetnet was generated from Hetionet v1.0, the second permuted hetnet was generated from the first, and so on. For each metaedge, we assessed the percent of edges unchanged as the algorithm progressed to ensure that a sufficient number of swaps had been performed to randomize the network [200]. Permuted hetnets are useful for computing the baseline performance of meaningless edges while preserving node degree [202]. Since, our use of permutation focused on assessing Δ AUROC, a small number of permuted hetnets was sufficient, as the variability in a metapath’s AUROC across the permuted hetnets was low.

### Graph databases & Neo4j

Traditional relational databases — such as SQLite, MySQL, and PostgreSQL — excel at storing highly structured data in tables. Connectivity between tables is accomplished using foreign-key references between columns. However, for many biomedical applications the connectivity between entities is of foremost importance. Furthermore, enforcing a rigid structure of what attributes an entity may possess is less important and often unnecessarily prohibitive. Graph databases focus instead on capturing connectivity (relationships) between entities (nodes). Accordingly, graph databases such as Neo4j offer greater ease when modeling biomedical relationships and superior performance when traversing many levels of connectivity [203,204]. Until recently, graph database adoption in bioinformatics was limited [205]. However lately, the demand to model and capture biological connectivity at scale has led to increasing adoption [206–209].

We used the Neo4j graph database for storing and operating on Hetionet and noticed major benefits from tapping into this large open source ecosystem [210]. Persistent storage with immediate access and the Cypher query language — a sort of SQL for hetnets — were two of the biggest benefits. To facilitate our migration to Neo4j, we updated hetio — our existing Python package for hetnets [211] — to export networks into Neo4j and DWPC queries to Cypher. In addition, we created an interactive GraphGist for Project Rephetio, which introduces our approach and showcases its Cypher queries. Finally, we created a public Neo4j instance [212], which leverages several modern technologies such Neo4j Browser guides, cloud hosting with HTTPS, and Docker deployment [213, 214].

### Machine learning approach

Project Rephetio relied on the previously-published DWPC metric to generate features for compound–disease pairs. The DWPC measures the prevalence of a given metapath between a given source and target node [22]. It is calculated by first extracting all paths from the source to target node that follow the specified metapath. Next, each path is weighted by taking the product of the node degrees along the path raised to a negative exponent. This damping exponent — the sole parameter — thereby determines the extent that paths through high-degree nodes are downweighted: we chose w = 0.4 based on our past optimizations [22]. The DWPC equals the sum of the path weights (referred to as path-degree products). Traversing the hetnet to extract all paths between a source and target node, which we performed in Neo4j, is the most computationally intensive step in computing DWPCs [215]. For future work, we are exploring matrix multiplication approaches, which could improve runtime several orders of magnitude.

Project Rephetio made several refinements to metapath-based hetnet edge prediction compared to previous studies [22,23]. First, we transformed DWPCs by mean scaling and then taking the inverse hyperbolic sine [216] to make them more amenable to modeling [217]. Second, we bifurcated the workflow into an all-features stage and an all-observations stage [40]. The all-features stage assesses feature performance and does not require computing features for all negatives. Here we selected a random subset of 3,020 (4 × 755) negatives. Little error was introduced by this optimization, since the predominant limitation to performance assessment was the small number of positives (755) rather than negatives. Based on the all-features performance assessment [218], we selected 142 DWPCs to compute on all observations (all 209,168 compound–disease pairs). The feature selection was designed to remove uninformative features (according to permutation) and guard against edge-dropout contamination [219]. Third, we included 14 degree features, which assess the degree of a specific metaedge for either the source compound or target disease.

### Network support of predictions

To improve the interpretability of the predictions, we developed a method for decomposing a prediction into its network support [220]. This information is deployed to our Neo4j Browser guides, allowing users to assess the biomedical evidence contributing to a given prediction. First, we used logistic regression terms to quantify the contribution of metapaths that positively support a prediction. Second, we decomposed a metapath’s contribution, according to its DWPC, into specific paths contributions. Finally, we aggregated paths based on their source (first) or target (last) edge to quantify the contribution of specific edges of the source compound or target disease [221].

Using the acamprosate–epilepsy prediction as an example, we first quantified metapath contributions: 40% of the prediction was supported by *CbGbCtD* paths, 36% by *CbGaD* paths, 11% by *CcSEcCtD* paths, 8% by *CbGpPWpGaD* paths, and 5% by *CbGeAlD* paths. Second, we calculated path contributions: *Acamprosate–binds–GRM5–associates–epilepsy* syndrome was the most supportive path, contributing 11% of the prediction. Finally, we aggregated path contributions to calculate that the source edge of *Acamprosate—binds—GRM5* contributed 23% of the prediction, while the target edge of *epilepsy syndrome–treats–Felbamate* contributed 12%.

### Prior probability of treatment

The 755 treatments in Hetionet v1.0 are not evenly distributed between all compounds and diseases. For example, methotrexate treats 19 diseases and hypertension is treated by 68 compounds. We estimated a prior probability of treatment — based only on the treatment degree of the source compound and target disease — on 744,975 permutations of the bipartite treatment network [222]. Methotrexate received a 79.6% prior probability of treating hypertension, whereas a compound and disease that both had only one treatment received a prior of 0.12%.

Across the 209,168 compound–disease pairs, the prior predicted the known treatments with AUROC = 97.9%. The strength of this association threatened to dominate our predictions. However, not modeling the prior can lead to omitted-variable bias and confounded proxy variables. To address the issue, we included the logit-transformed prior, without any regularization, as a term in the model. This restricted model fitting to the 29,799 observations with a nonzero prior — corresponding to the 387 compounds and 77 diseases with at least one treatment. To enable predictions for all 209,168 observations, we set the prior for each compound–disease pair to the overall prevalence of positives (0.36%).

This method succeeded at accommodating the treatment degrees. The prior probabilities performed poorly on the validation sets with AUROC = 54.1% on DrugCentral indications and AUROC = 62.5% on clinical trials. This performance dropoff compared to training shows the danger of encoding treatment degree into predictions. The benefits of our solution are highlighted by the superior validation performance of our predictions compared to the prior (Figure 3).

### Indication sets

We evaluated our predictions on four sets of indications as shown in Figure 3.

- **Disease Modifying**— the 755 disease modifying treatments in PharmacotherapyDB v1.0. These indications are included in the hetnet as treats edges and used to train the logistic regression model. Due to edge dropout contamination and self-testing [219, 220], overfitting could potentially inflate performance on this set. Therefore, for the three remaining indication sets, we removed any observations that were positives in this set.
- **DrugCentral**— We discovered the DrugCentral database after completing our physician curation for PharmacotherapyDB. This database contained 210 additional indications [92]. While we didn’t curate these indications, we observed a high proportion of disease modifying therapy.
- **Clinical Trial**— We compiled indications that have been investigated by clinical trial from ClinicalTrials.gov [224]. This set contains 5,594 indications. Since these indications were not manually curated and clinical trials often show a lack of efficacy, we expected lower performance on this set.
- **Symptomatic** — 390 symptomatic indications from PharacotherapyDB. These edges are included in the hetnet as *palliates* edges.

Only the Clinical Trial and DrugCentral indication sets were used for external validation, since the Disease Modifying and Symptomatic indications were included in the hetnet. As an aside, several additional indication catalogs have recently been published, which future studies may want to also consider [173,225–227].

### Realtime open science & Thinklab

We conducted our study using Thinklab — a platform for realtime open collaborative science — on which this study was the first project. We began the study by publicly proposing the idea and inviting discussion [228]. We continued by chronicling our progress via discussions. We used Thinklab as the frontend to coordinate and report our analyses and GitHub as the backend to host our code, data, and notebooks. On top of our Thinklab team consisting of core contributors, we welcomed community contribution and review. In areas where our expertise was lacking or advice would be helpful, we sought input from domain experts and encouraged them to respond on Thinklab where their comments would be CC BY licensed and their contribution rated and rewarded.

In total, 40 non-team members commented across 86 discussions, which generated 622 comments and 191 notes (Figure 6). Thinklab content for this project totaled 145,771 words or 918,837 characters [229]. Using an estimated 7,000 words per academic publication as a benchmark, Project Rephetio generated written content comparable in volume to 20.8 publications prior to its completion. We noticed several other benefits from using Thinklab including forging a community of contributors [230]; receiving feedback during the early stages when feedback was most actionable [231]; disseminating our research without delay [232,233]; opening avenues for external input [234]; facilitating problem-oriented teaching [235,236]; and improving our documentation by maintaining a publication-grade digital lab notebook [237].

**Figure 6:**
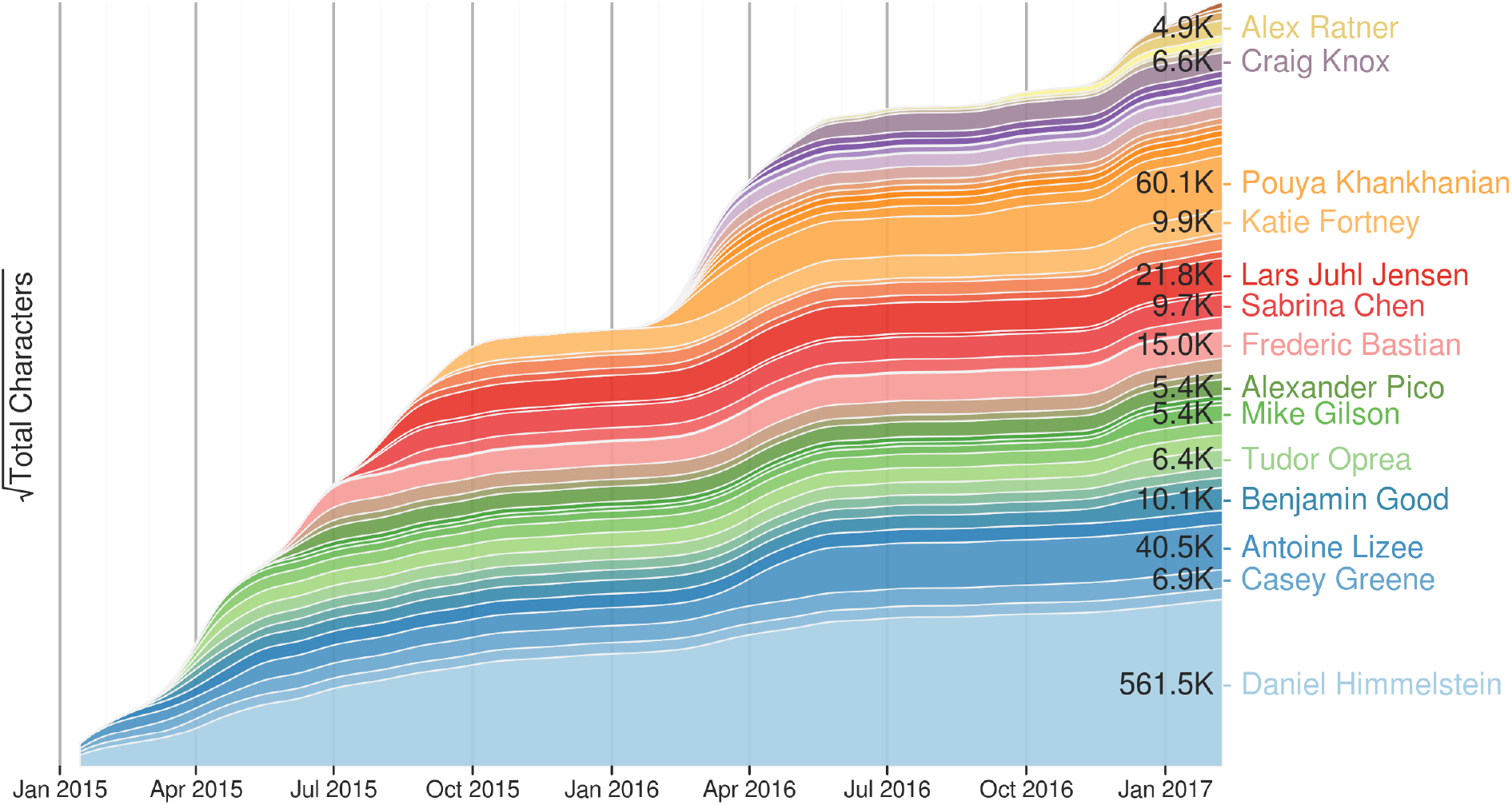
The growth the Project Rephetio corpus on Thinklab over time. This figure shows Project Rephetio contributions by user over time. Each band represented the cumulative contribution of a Thinklab user to discussions in the Rephetio project [229]. Users are ordered by date of first contribution. Users who contributed over 4,500 characters are named. The square root transformation of characters written per user accentuates the activity of new contributors, thereby emphasizing collaboration and diverse input. Thinklab began winding down operations in July 2017 and has switched to a static state. While users will no longer be able to add comments, the corpus of content remains browsable at https://think-lab.github.io and available in machine-readable formats at https://github.com/dhimmel/thinklytics.

## Acknowledgements

We are immensely grateful to our Thinklab contributors who joined us in our experiment of radically open science. The following non-team members provided contributions that received 5 or more Thinklab points: Lars Juhl Jensen, Frederic Bastian, Alexander Pico, Casey Greene, Benjamin Good, Craig Knox, Mike Gilson, Chris Mungall, Katie Fortney, Venkat Malladi, Tudor Oprea, MacKenzie Smith, Caty Chung, Allison McCoy, Alexey Strokach, Ritu Khare, Greg Way, Marina Sirota, Raghavendran Partha, Oleg Ursu, Jesse Spaulding, Gaya Nadarajan, Alex Ratner, Scooter Morris, Alessandro Didonna, Alex Pankov, Tong Shu Li, and Janet Piñero. Additionally, the founder of Thinklab, Jesse Spaulding, supported community contributions and developed the platform with Project Rephetio’s needs in mind. We also appreciate DigitalOcean’s sponsorship the Hetionet Browser to cover its hosting costs. Finally, we would like to thank Neo Technology, whose staff provided excellent technical support.

This material is based upon work supported by the National Science Foundation Graduate Research Fellowship under Grant No. 1144247 to DSH. SEB is supported by the Heidrich Family and Friends Foundation.

## References

1. DiMasi JA, Grabowski HG, Hansen RW. 2016 Innovation in the pharmaceutical industry: New estimates of R&D costs. Journal of Health Economics 47, 20–33. See https://doi.org/10.1016/j.jhealeco.2016.01.012.

2. Reichert JM. 2003 A guide to drug discovery: Trends in development and approval times for new therapeutics in the United States. Nat Rev Drug Discov 2, 695–702. See https://doi.org/10.1038/nrd1178.

3. Hay M, Thomas DW, Craighead JL, Economides C, Rosenthal J. 2014 Clinical development success rates for investigational drugs. Nat Biotechnol 32, 40–51. See https://doi.org/10.1038/nbt.2786.

4. Scannell JW, Blanckley A, Boldon H, Warrington B. 2012 Diagnosing the decline in pharmaceutical R&D efficiency. Nat Rev Drug Discov 11, 191–200. See https://doi.org/10.1038/nrd3681.

5. Ashburn TT, Thor KB. 2004 Drug repositioning: identifying and developing new uses for existing drugs. Nat Rev Drug Discov 3, 673–683. See https://doi.org/10.1038/nrd1468.

6. Wang G, Jung K, Winnenburg R, Shah NH. 2015 A method for systematic discovery of adverse drug events from clinical notes. Journal of the American Medical Informatics Association 22, 1196–1204. See https://doi.org/10.1093/jamia/ocv102.

7. Xu H et al. 2014 Validating drug repurposing signals using electronic health records: a case study of metformin associated with reduced cancer mortality. Journal of the American Medical Informatics Association See https://doi.org/10.1136/amiajnl-2014-002649.

8. Brilliant MH et al. 2016 Mining Retrospective Data for Virtual Prospective Drug Repurposing: L-DOPA and Age-related Macular Degeneration. The American Journal of Medicine 129, 292–298. See https://doi.org/10.1016/j.amjmed.2015.10.015.

9. Tatonetti NP, Ye PP, Daneshjou R, Altman RB. 2012 Data-Driven Prediction of Drug Effects and Interactions. Science Translational Medicine 4, 125ra31–125ra31. See https://doi.org/10.1126/scitranslmed.3003377.

10. Stephens M, Balding DJ. 2009 Bayesian statistical methods for genetic association studies. Nat Rev Genet 10, 681–690. See https://doi.org/10.1038/nrg2615.

11. Sawcer S. 2008 The complex genetics of multiple sclerosis: pitfalls and prospects. Brain 131, 3118–3131. See https://doi.org/10.1093/brain/awn081.

12. Roth BL, Sheffler DJ, Kroeze WK. 2004 Magic shotguns versus magic bullets: selectively nonselective drugs for mood disorders and schizophrenia. Nat Rev Drug Discov 3, 353–359. See https://doi.org/10.1038/nrd1346.

13. Hopkins AL. 2008 Network pharmacology: the next paradigm in drug discovery. Nat Chem Biol 4, 682–690. See https://doi.org/10.1038/nchembio.118.

14. Hopkins AL. 2007 Network pharmacology. Nat Biotechnol 25, 1110–1111. See https://doi.org/10.1038/nbt1007-1110.

15. Swinney DC, Anthony J. 2011 How were new medicines discovered? Nat Rev Drug Discov 10, 507–519. See https://doi.org/10.1038/nrd3480.

16. Iskar M, Zeller G, Zhao X-M, van Noort V, Bork P. 2012 Drug discovery in the age of systems biology: the rise of computational approaches for data integration. Current Opinion in *Biotechnology* 23, 609–616. See https://doi.org/10.1016/j.copbio.2011.11.010.

17. Lamb J. 2007 The Connectivity Map: a new tool for biomedical research. Nat Rev Cancer 7, 54–60. See https://doi.org/10.1038/nrc2044.

18. Qu XA, Rajpal DK. 2012 Applications of Connectivity Map in drug discovery and development. Drug Discovery Today 17, 1289–1298. See https://doi.org/10.1016/j.drudis.2012.07.017.

19. Hodos RA, Kidd BA, Shameer K, Readhead BP, Dudley JT. 2016 In silicomethods for drug repurposing and pharmacology. WIREs Syst Biol Med 8, 186–210. See https://doi.org/10.1002/wsbm.1337.

20. Hurle MR, Yang L, Xie Q, Rajpal DK, Sanseau P, Agarwal P. 2013 Computational Drug Repositioning: From Data to Therapeutics. Clin Pharmacol Ther 93, 335–341. See https://doi.org/10.1038/clpt.2013.1.

21. Liu Z, Fang H, Reagan K, Xu X, Mendrick DL, Slikker W Jr, Tong W. 2013 In silico drug repositioning – what we need to know. Drug Discovery Today 18, 110–115. See https://doi.org/10.1016/j.drudis.2012.08.005.

22. Himmelstein DS, Baranzini SE. 2015 Heterogeneous Network Edge Prediction: A Data Integration Approach to Prioritize Disease-Associated Genes. PLoS Comput Biol 11, e1004259. See https://doi.org/10.1371/journal.pcbi.1004259.

23. Sun Y, Barber R, Gupta M, Aggarwal CC, Han J. 2011 Co-author Relationship Prediction in Heterogeneous Bibliographic Networks. In 2011 International Conference on Advances in Social Networks Analysis and Mining, IEEE. See https://doi.org/10.1109/asonam.2011.112.

24. Gottlieb A, Stein GY, Ruppin E, Sharan R. 2014 PREDICT: a method for inferring novel drug indications with application to personalized medicine. Molecular Systems Biology 7, 496–496. See https://doi.org/10.1038/msb.2011.26.

25. Cheng J, Yang L, Kumar V, Agarwal P. 2014 Systematic evaluation of connectivity map for disease indications. Genome Med 6. See https://doi.org/10.1186/s13073-014-0095-1.

26. Guney E, Menche J, Vidal M, Barábasi A-L. 2016 Network-based in silico drug efficacy screening. Nat Comms 7, 10331. See https://doi.org/10.1038/ncomms10331.

27. Li J, Lu Z. 2012 A new method for computational drug repositioning using drug pairwise similarity. In 2012 IEEE International Conference on Bioinformatics and Biomedicine, IEEE. See https://doi.org/10.1109/bibm.2012.6392722.

28. Chiang AP, Butte AJ. 2009 Systematic Evaluation of Drug–Disease Relationships to Identify Leads for Novel Drug Uses. Clin Pharmacol Ther 86, 507–510. See https://doi.org/10.1038/clpt.2009.103.

29. Lamb J. 2006 The Connectivity Map: Using Gene-Expression Signatures to Connect Small Molecules, Genes, and Disease. Science 313, 1929–1935. See https://doi.org/10.1126/science.1132939.

30. Iorio F, Rittman T, Ge H, Menden M, Saez-Rodriguez J. 2013 Transcriptional data: a new gateway to drug repositioning? Drug Discovery Today 18, 350–357. See https://doi.org/10.1016/j.drudis.2012.07.014.

31. Nelson MR et al. 2015 The support of human genetic evidence for approved drug indications. Nat Genet 47, 856–860. See https://doi.org/10.1038/ng.3314.

32. Sanseau P, Agarwal P, Barnes MR, Pastinen T, Richards JB, Cardon LR, Mooser V. 2012 Use of genome-wide association studies for drug repositioning. Nat Biotechnol 30, 317–320. See https://doi.org/10.1038/nbt.2151.

33. Campillos M, Kuhn M, Gavin A-C, Jensen LJ, Bork P. 2008 Drug Target Identification Using Side-Effect Similarity. Science 321, 263–266. See https://doi.org/10.1126/science.1158140.

34. Nugent T, Plachouras V, Leidner JL. 2016 Computational drug repositioning based on side-effects mined from social media. PeerJ Computer Science 2, e46. See https://doi.org/10.7717/peerj-cs.46.

35. Zhou X, Menche J, Barabási A-L, Sharma A. 2014 Human symptoms–disease network. Nat Comms 5. See https://doi.org/10.1038/ncomms5212.

36. Pratanwanich N, Lió P. 2014 Pathway-based Bayesian inference of drug–disease interactions. Mol. BioSyst. 10, 1538–1548. See https://doi.org/10.1039/c4mb00014e.

37. Himmelstein D. 2016 Exploring the power of Hetionet: a Cypher query depot. See https://doi.org/10.15363/thinklab.d220.

38. Himmelstein D. 2017 Dhimmel/Hetionet V1.0.0: Hetionet V1.0 In Json, Tsv, And Neo4J Formats. See https://doi.org/10.5281/zenodo.268568.

39. Himmelstein D, Lizee A. 2016 Computing standardized logistic regression coefficients. See https://doi.org/10.15363/thinklab.d205.

40. Himmelstein D. 2016 Our hetnet edge prediction methodology: the modeling framework for Project Rephetio. See https://doi.org/10.15363/thinklab.d210.

41. Himmelstein D. 2017 Dhimmel/Learn V1.0: The Machine Learning Repository For Project Rephetio. See https://doi.org/10.5281/zenodo.268654.

42. Himmelstein D, Hessler C, Khankhanian P. 2016 Predictions of whether a compound treats a disease. See https://doi.org/10.15363/thinklab.d203.

43. Harmey D, Griffin PR, Kenny PJ. 2012 Development of Novel Pharmacotherapeutics for Tobacco Dependence: Progress and Future Directions. Nicotine & Tobacco Research 14, 1300–1318. See https://doi.org/10.1093/ntr/nts201.

44. Mihalak KB. 2006 Varenicline Is a Partial Agonist at 4beta2 and a Full Agonist at 7 Neuronal Nicotinic Receptors. Molecular Pharmacology 70, 801–805. See https://doi.org/10.1124/mol.106.025130.

45. Thorgeirsson TE et al. 2008 A variant associated with nicotine dependence, lung cancer and peripheral arterial disease. Nature 452, 638–642. See https://doi.org/10.1038/nature06846.

46. Boshier A, Wilton LV, Shakir SAW. 2003 Evaluation of the safety of bupropion (Zyban) for smoking cessation from experience gained in general practice use in England in 2000. European Journal of Clinical Pharmacology 59, 767–773. See https://doi.org/10.1007/s00228-003-0693-0.

47. Hays JT, Ebbert JO, Sood A. 2008 Efficacy and Safety of Varenicline for Smoking Cessation. The American Journal of Medicine 121, S32–S42. See https://doi.org/10.1016/j.amjmed.2008.01.017.

48. Cahill K, Lindson-Hawley N, Thomas KH, Fanshawe TR, Lancaster T. 2016 Nicotine receptor partial agonists for smoking cessation. Cochrane Database of Systematic Reviews. See https://doi.org/10.1002/14651858.cd006103.pub7.

49. West R, Zatonski W, Cedzynska M, Lewandowska D, Pazik J, Aveyard P, Stapleton J. 2011 Placebo-Controlled Trial of Cytisine for Smoking Cessation. N Engl J Med 365, 1193–1200. See https://doi.org/10.1056/nejmoa1102035.

50. Walker N, Howe C, Glover M, McRobbie H, Barnes J, Nosa V, Parag V, Bassett B, Bullen C. 2014 Cytisine versus Nicotine for Smoking Cessation. N Engl J Med 371, 2353–2362. See https://doi.org/10.1056/nejmoa1407764.

51. Ashare RL, Kimmey BA, Rupprecht LE, Bowers ME, Hayes MR, Schmidt HD. 2016 Repeated administration of an acetylcholinesterase inhibitor attenuates nicotine taking in rats and smoking behavior in human smokers. Transl Psychiatry 6, e713. See https://doi.org/10.1038/tp.2015.209.

52. Khankhanian P, Himmelstein D. 2016 Prediction in epilepsy. See https://doi.org/10.15363/thinklab.d224.

53. Himmelstein D, Khankhanian P, Pico A, Jensen LJ, Morris S. 2017 Visualizing the top epilepsy predictions in Cytoscape. See https://doi.org/10.15363/thinklab.d230.

54. Mirsattari SM, Sharpe MD, Young GB. 2004 Treatment of Refractory Status Epilepticus With Inhalational Anesthetic Agents Isoflurane and Desflurane. Arch Neurol 61. See https://doi.org/10.1001/archneur.61.8.1254.

55. In press. Anatomical Therapeutic Chemical Classification System (WHO). The SAGE Encyclopedia of Pharmacology and Society. See https://doi.org/10.4135/9781483349985.n37.

56. I. Johannessen S, Johannessen Landmark C. 2010 Antiepileptic Drug Interactions - Principles and Clinical Implications. CN 8, 254–267. See https://doi.org/10.2174/157015910792246254.

57. Rogawski MA, Löscher W. 2004 The neurobiology of antiepileptic drugs. Nat Rev Neurosci 5, 553–564. See https://doi.org/10.1038/nrn1430.

58. Johannessen Landmark C, Henning O, Johannessen SI. 2016 Proconvulsant effects of antidepressants — What is the current evidence? Epilepsy & Behavior 61, 287–291. See https://doi.org/10.1016/j.yebeh.2016.01.029.

59. Himmelstein D. 2017 Why we predicted ictogenic tricyclic compounds treat epilepsy? See https://doi.org/10.15363/thinklab.d231.

60. Dailey JW, Naritoku DK. 1996 Antidepressants and seizures: Clinical anecdotes overshadow neuroscience. Biochemical Pharmacology 52, 1323–1329. See https://doi.org/10.1016/s0006-2952(96)00509-6.

61. Zadikoff C, Munhoz RP, Asante AN, Politzer N, Wennberg R, Carlen P, Lang A. 2007 Movement disorders in patients taking anticonvulsants. Journal of Neurology, Neurosurgery & Psychiatry 78, 147–151. See https://doi.org/10.1136/jnnp.2006.100222.

62. Wu D, Thijs RD. 2015 Anticonvulsant-induced downbeat nystagmus in epilepsy. Epilepsy & Behavior Case Reports 4, 74–75. See https://doi.org/10.1016/j.ebcr.2015.07.003.

63. Roff Hilton EJ, Hosking SL, Betts T. 2004 The effect of antiepileptic drugs on visual performance. Seizure 13, 113–128. See https://doi.org/10.1016/s1059-1311(03)00082-7.

64. Placidi F, Scalise A, Marciani MG, Romigi A, Diomedi M, Gigli GL. 2000 Effect of antiepileptic drugs on sleep. Clinical Neurophysiology 111, S115–S119. See https://doi.org/10.1016/s1388-2457(00)00411-9.

65. Jahromi SR, Togha M, Fesharaki SH, Najafi M, Moghadam NB, Kheradmand JA, Kazemi H, Gorji A. 2011 Gastrointestinal adverse effects of antiepileptic drugs in intractable epileptic patients. Seizure 20, 343–346. See https://doi.org/10.1016/j.seizure.2010.12.011.

66. Gligorijević V, Pržulj N. 2015 Methods for biological data integration: perspectives and challenges. J. R. Soc. Interface 12, 20150571. See https://doi.org/10.1098/rsif.2015.0571.

67. Kivela M, Arenas A, Barthelemy M, Gleeson JP, Moreno Y, Porter MA. 2014 Multilayer networks. Journal of Complex Networks 2, 203–271. See https://doi.org/10.1093/comnet/cnu016.

68. Himmelstein D, Greene C, Baranzini S. 2015 Renaming ‘heterogeneous networks’ to a more concise and catchy term. See https://doi.org/10.15363/thinklab.d104.

69. Himmelstein D, Lizee A, Hessler C, Brueggeman L, Chen S, Hadley D, Green A, Khankhanian P, Baranzini S. 2015 Rephetio: Repurposing drugs on a hetnet [project]. See https://doi.org/10.15363/thinklab.4.

70. Sirota M, Dudley JT, Kim J, Chiang AP, Morgan AA, Sweet-Cordero A, Sage J, Butte AJ. 2011 Discovery and Preclinical Validation of Drug Indications Using Compendia of Public Gene Expression Data. Science Translational Medicine 3, 96ra77–96ra77. See https://doi.org/10.1126/scitranslmed.3001318.

71. Farook JM, Krazem A, Lewis B, Morrell DJ, Littleton JM, Barron S. 2008 Acamprosate attenuates the handling induced convulsions during alcohol withdrawal in Swiss Webster mice. Physiology & Behavior 95, 267–270. See https://doi.org/10.1016/j.physbeh.2008.05.020.

72. Ehrenberg HR, Shin J, Ratner AJ, Fries JA, Ré C. 2016 Data programming with DDLite. In Proceedings of the Workshop on Human-In-the-Loop Data Analytics - HILDA ‘16, ACM Press. See https://doi.org/10.1145/2939502.2939515.

73. Himmelstein D, Good B, Khankhanian P, Ratner A. 2016 Brainstorming future directions for Hetionet. See https://doi.org/10.15363/thinklab.d227.

74. Chen PP-S. 1997 English, Chinese and ER diagrams. Data & Knowledge Engineering 23, 5–16. See https://doi.org/10.1016/s0169-023x(97)00017-7.

75. Himmelstein D, Jensen LJ, Khankhanian P. 2016 Data nomenclature: naming and abbreviating our network types. See https://doi.org/10.15363/thinklab.d162.

76. Malone J, Stevens R, Jupp S, Hancocks T, Parkinson H, Brooksbank C. 2016 Ten Simple Rules for Selecting a Bio-ontology. PLoS Comput Biol 12, e1004743. See https://doi.org/10.1371/journal.pcbi.1004743.

77. Schriml LM, Arze C, Nadendla S, Chang Y-WW, Mazaitis M, Felix V, Feng G, Kibbe WA. 2011 Disease Ontology: a backbone for disease semantic integration. Nucleic Acids Research 40, D940–D946. See https://doi.org/10.1093/nar/gkr972.

78. Kibbe WA et al. 2014 Disease Ontology 2015 update: an expanded and updated database of human diseases for linking biomedical knowledge through disease data. Nucleic Acids Research 43, D1071–D1078. See https://doi.org/10.1093/nar/gku1011.

79. Himmelstein D, Li TS. 2015 Unifying disease vocabularies. See https://doi.org/10.15363/thinklab.d44.

80. Himmelstein DS. 2016 User-Friendly Extensions To The Disease Ontology V1.0. See https://doi.org/10.5281/zenodo.45584.

81. Wu T-J et al. 2015 Generating a focused view of disease ontology cancer terms for pan-cancer data integration and analysis. Database 2015, bav032–bav032. See https://doi.org/10.1093/database/bav032.

82. Himmelstein D, Pankov A. 2015 Mining knowledge from MEDLINE articles and their indexed MeSH terms. See https://doi.org/10.15363/thinklab.d67.

83. Himmelstein DS. 2016 User-Friendly Extensions To Mesh V1.0. See https://doi.org/10.5281/zenodo.45586.

84. Law V et al. 2013 DrugBank 4.0: shedding new light on drug metabolism. Nucl. Acids Res. 42, D1091–D1097. See https://doi.org/10.1093/nar/gkt1068.

85. Himmelstein D. 2015 Unifying drug vocabularies. See https://doi.org/10.15363/thinklab.d40.

86. Himmelstein DS. 2016 User-Friendly Extensions Of The Drugbank Database V1.0. See https://doi.org/10.5281/zenodo.45579.

87. Kuhn M, Letunic I, Jensen LJ, Bork P. 2015 The SIDER database of drugs and side effects. Nucleic Acids Res 44, D1075–D1079. See https://doi.org/10.1093/nar/gkv1075.

88. Himmelstein D. 2015 Extracting side effects from SIDER 4. See https://doi.org/10.15363/thinklab.d97.

89. Himmelstein DS. 2016 Extracting Tidy And User-Friendly Tsvs From Sider 4.1. See https://doi.org/10.5281/zenodo.45521.

90. Bodenreider O. 2004 The Unified Medical Language System (UMLS): integrating biomedical terminology. Nucleic Acids Research 32, 267D–270. See https://doi.org/10.1093/nar/gkh061.

91. Ursu O, Holmes J, Knockel J, Bologa CG, Yang JJ, Mathias SL, Nelson SJ, Oprea TI. 2016 DrugCentral: online drug compendium. Nucleic Acids Res 45, D932–D939. See https://doi.org/10.1093/nar/gkw993.

92. Himmelstein D, Ursu O, Gilson M, Khankhanian P, Oprea T. 2016 Incorporating DrugCentral data in our network. See https://doi.org/10.15363/thinklab.d186.

93. Maglott D, Ostell J, Pruitt KD, Tatusova T. 2010 Entrez Gene: gene-centered information at NCBI. Nucleic Acids Research 39, D52–D57. See https://doi.org/10.1093/nar/gkq1237.

94. Himmelstein D, Greene C, Pico A. 2015 Using Entrez Gene as our gene vocabulary. See https://doi.org/10.15363/thinklab.d34.

95. Himmelstein DS. 2016 Processed Entrez Gene Datasets For Humans V1.0. See https://doi.org/10.5281/zenodo.45524.

96. Mungall CJ, Torniai C, Gkoutos GV, Lewis SE, Haendel MA. 2012 Uberon, an integrative multispecies anatomy ontology. Genome Biol 13, R5. See https://doi.org/10.1186/gb-2012-13-1-r5.

97. Malladi V, Himmelstein D, Mungall C. 2015 Tissue Node. See https://doi.org/10.15363/thinklab.d41.

98. Himmelstein DS. 2016 User-Friendly Anatomical Structures Data From The Uberon Ontology V1.0. See https://doi.org/10.5281/zenodo.45527.

99. Kutmon M et al. 2015 WikiPathways: capturing the full diversity of pathway knowledge. Nucleic Acids Res 44, D488–D494. See https://doi.org/10.1093/nar/gkv1024.

100. Pico AR, Kelder T, van Iersel MP, Hanspers K, Conklin BR, Evelo C. 2008 WikiPathways: Pathway Editing for the People. PLoS Biol 6, e184. See https://doi.org/10.1371/journal.pbio.0060184.

101. Fabregat A et al. 2015 The Reactome pathway Knowledgebase. Nucleic Acids Res 44, D481–D487. See https://doi.org/10.1093/nar/gkv1351.

102. Schaefer CF, Anthony K, Krupa S, Buchoff J, Day M, Hannay T, Buetow KH. 2008 PID: the Pathway Interaction Database. Nucleic Acids Research 37, D674–D679. See https://doi.org/10.1093/nar/gkn653.

103. Cerami EG, Gross BE, Demir E, Rodchenkov I, Babur O, Anwar N, Schultz N, Bader GD, Sander C. 2010 Pathway Commons, a web resource for biological pathway data. Nucleic Acids Research 39, D685–D690. See https://doi.org/10.1093/nar/gkq1039.

104. Pico A, Himmelstein D. 2015 Adding pathway resources to your network. See https://doi.org/10.15363/thinklab.d72.

105. Himmelstein DS, Pico AR. 2016 Dhimmel/Pathways V2.0: Compiling Human Pathway Gene Sets. See https://doi.org/10.5281/zenodo.48810.

106. Ashburner M et al. 2000 Gene Ontology: tool for the unification of biology. Nat Genet 25, 25–29. See https://doi.org/10.1038/75556.

107. Himmelstein D. 2015 Disease Ontology feature requests. See https://doi.org/10.15363/thinklab.d68.

108. Hersey A, Chambers J, Bellis L, Patrícia Bento A, Gaulton A, Overington JP. 2015 Chemical databases: curation or integration by user-defined equivalence? Drug Discovery Today: Technologies 14, 17–24. See https://doi.org/10.1016/j.ddtec.2015.01.005.

109. Chambers J et al. 2013 UniChem: a unified chemical structure cross-referencing and identifier tracking system. Journal of Cheminformatics 5, 3. See https://doi.org/10.1186/1758-2946-5-3.

110. Chambers J, Davies M, Gaulton A, Papadatos G, Hersey A, Overington JP. 2014 UniChem: extension of InChI-based compound mapping to salt, connectivity and stereochemistry layers. J Cheminform 6. See https://doi.org/10.1186/s13321-014-0043-5.

111. Heller S, McNaught A, Stein S, Tchekhovskoi D, Pletnev I. 2013 InChI - the worldwide chemical structure identifier standard. Journal of Cheminformatics 5, 7. See https://doi.org/10.1186/1758-2946-5-7.

112. Himmelstein D, Bastian F, Baranzini S. 2016 Dhimmel/Bgee V1.0: Anatomy-Specific Gene Expression In Humans From Bgee. See https://doi.org/10.5281/zenodo.47157.

113. Himmelstein D, Bastian F. 2015 Processing Bgee for tissue-specific gene presence and over/under-expression. See https://doi.org/10.15363/thinklab.d124.

114. Himmelstein D, Bastian F. 2015 Tissue-specific gene expression resources. See https://doi.org/10.15363/thinklab.d81.

115. Bastian F, Parmentier G, Roux J, Moretti S, Laudet V, Robinson-Rechavi M. In press. Bgee: Integrating and Comparing Heterogeneous Transcriptome Data Among Species. In Lecture Notes in Computer Science, pp. 124–131. Springer Berlin Heidelberg. See https://doi.org/10.1007/978-3-540-69828-9_12.

116. Santos A, Tsafou K, Stolte C, Pletscher-Frankild S, O’Donoghue SI, Jensen LJ. 2015 Comprehensive comparison of large-scale tissue expression datasets. PeerJ 3, e1054. See https://doi.org/10.7717/peerj.1054.

117. Himmelstein D, Jensen LJ. 2015 Gene–Tissue Relationships From The Tissues Database. See https://doi.org/10.5281/zenodo.27244.

118. Himmelstein D, Jensen LJ. 2015 The TISSUES resource for the tissue-specificity of genes. See https://doi.org/10.15363/thinklab.d91.

119. Chen X, Liu M, Gilson M. 2001 BindingDB: A Web-Accessible Molecular Recognition Database. CCHTS 4, 719–725. See https://doi.org/10.2174/1386207013330670.

120. Gilson MK, Liu T, Baitaluk M, Nicola G, Hwang L, Chong J. 2015 BindingDB in 2015: A public database for medicinal chemistry, computational chemistry and systems pharmacology. Nucleic Acids Res 44, D1045–D1053. See https://doi.org/10.1093/nar/gkv1072.

121. Wishart DS. 2006 DrugBank: a comprehensive resource for in silico drug discovery and exploration. Nucleic Acids Research 34, D668–D672. See https://doi.org/10.1093/nar/gkj067.

122. Himmelstein D, Gilson M. 2015 Integrating drug target information from BindingDB. See https://doi.org/10.15363/thinklab.d53.

123. Himmelstein D, Gilson M, Baranzini S. 2015 Processing The October 2015 Bindingdb. See https://doi.org/10.5281/zenodo.33987.

124. Himmelstein D, Chen S. 2015 Protein (target, carrier, transporter, and enzyme) interactions in DrugBank. See https://doi.org/10.15363/thinklab.d65.

125. Himmelstein D, Chen S. 2015 Calculating molecular similarities between DrugBank compounds. See https://doi.org/10.15363/thinklab.d70.

126. Himmelstein D, Brueggeman L, Baranzini S. 2015 Pairwise molecular similarities between DrugBank compounds. See https://doi.org/10.6084/m9.figshare.1418386.

127. Dice LR. 1945 Measures of the Amount of Ecologic Association Between Species. Ecology 26, 297–302. See https://doi.org/10.2307/1932409.

128. Rogers D, Hahn M. 2010 Extended-Connectivity Fingerprints. J. Chem. Inf. Model. 50, 742–754. See https://doi.org/10.1021/ci100050t.

129. Morgan HL. 1965 The Generation of a Unique Machine Description for Chemical Structures-A Technique Developed at Chemical Abstracts Service. J. Chem. Doc. 5, 107–113. See https://doi.org/10.1021/c160017a018.

130. Himmelstein DS, Baranzini SE. 2016 Dhimmel/Gwas-Catalog V1.0: Extracting Gene–Disease Associations From The Gwas Catalog. See https://doi.org/10.5281/zenodo.48428.

131. Himmelstein D, Jensen LJ. 2015 Processing the DISEASES resource for disease–gene relationships. See https://doi.org/10.15363/thinklab.d106.

132. Himmelstein DS, Jensen LJ. 2016 Dhimmel/Diseases V1.0: Processing The Diseases Database Of Gene–Disease Associations. See https://doi.org/10.5281/zenodo.48425.

133. Himmelstein D, piñero janet. 2015 Processing DisGeNET for disease-gene relationships. See https://doi.org/10.15363/thinklab.d105.

134. Himmelstein DS, Piñero J. 2016 Dhimmel/Disgenet V1.0: Processing The Disgenet Database Of Gene–Disease Associations. See https://doi.org/10.5281/zenodo.48426.

135. Himmelstein D. 2015 Functional disease annotations for genes using DOAF. See https://doi.org/10.15363/thinklab.d94.

136. Himmelstein DS. 2016 Dhimmel/Doaf V1.0: Processing The Doaf Database Of Gene–Disease Associations. See https://doi.org/10.5281/zenodo.48427.

137. MacArthur J et al. 2016 The new NHGRI-EBI Catalog of published genome-wide association studies (GWAS Catalog). Nucleic Acids Res 45, D896–D901. See https://doi.org/10.1093/nar/gkw1133.

138. Himmelstein D. 2015 Extracting disease-gene associations from the GWAS Catalog. See https://doi.org/10.15363/thinklab.d80.

139. Himmelstein D, Sirota M, Way G. 2015 Calculating genomic windows for GWAS lead SNPs. See https://doi.org/10.15363/thinklab.d71.

140. Pletscher-Frankild S, Pallejà A, Tsafou K, Binder JX, Jensen LJ. 2015 DISEASES: Text mining and data integration of disease–gene associations. Methods 74, 83–89. See https://doi.org/10.1016/j.ymeth.2014.11.020.

141. Pinero J, Queralt-Rosinach N, Bravo A, Deu-Pons J, Bauer-Mehren A, Baron M, Sanz F, Furlong LI. 2015 DisGeNET: a discovery platform for the dynamical exploration of human diseases and their genes. Database 2015, bav028–bav028. See https://doi.org/10.1093/database/bav028.

142. Piñero J, Bravo À, Queralt-Rosinach N, Gutiérrez-Sacristán A, Deu-Pons J, Centeno E, García-García J, Sanz F, Furlong LI. 2016 DisGeNET: a comprehensive platform integrating information on human disease-associated genes and variants. Nucleic Acids Res 45, D833–D839. See https://doi.org/10.1093/nar/gkw943.

143. Xu W, Wang H, Cheng W, Fu D, Xia T, Kibbe WA, Lin SM. 2012 A Framework for Annotating Human Genome in Disease Context. PLoS ONE 7, e49686. See https://doi.org/10.1371/journal.pone.0049686.

144. Himmelstein D, Bastian F, Hadley D, Greene C. 2015 STARGEO: expression signatures for disease using crowdsourced GEO annotation. See https://doi.org/10.15363/thinklab.d96.

145. Himmelstein D, Hadley D, Schepanovski A. 2016 Dhimmel/Stargeo V1.0: Differentially Expressed Genes For 48 Diseases From Stargeo. See https://doi.org/10.5281/zenodo.46866.

146. Himmelstein DS. 2016 Dhimmel/Medline V1.0: Disease, Symptom, And Anatomy Cooccurence In Medline. See https://doi.org/10.5281/zenodo.48445.

147. Himmelstein D. 2015 Disease similarity from MEDLINE topic cooccurrence. See https://doi.org/10.15363/thinklab.d93.

148. Fisher RA. 1922 On the Interpretation of χ 2 from Contingency Tables, and the Calculation of P. Journal of the Royal Statistical Society 85, 87. See https://doi.org/10.2307/2340521.

149. Himmelstein D, Chung C. 2015 Computing consensus transcriptional profiles for LINCS L1000 perturbations. See https://doi.org/10.15363/thinklab.d43.

150. Himmelstein D, Brueggeman L, Baranzini S. 2016 Consensus signatures for LINCS L1000 perturbations. See https://doi.org/10.6084/m9.figshare.3085426.v1.

151. Priedigkeit N, Wolfe N, Clark NL. 2015 Evolutionary Signatures amongst Disease Genes Permit Novel Methods for Gene Prioritization and Construction of Informative Gene-Based Networks. PLoS Genet 11, e1004967. See https://doi.org/10.1371/journal.pgen.1004967.

152. Himmelstein D, Partha R. 2015 Selecting informative ERC (evolutionary rate covariation) values between genes. See https://doi.org/10.15363/thinklab.d57.

153. Himmelstein DS. 2016 Dhimmel/Erc V1.0: Processing Human Evolutionary Rate Covaration Data. See https://doi.org/10.5281/zenodo.48444.

154. Himmelstein D, Hadley D, Strokach A. 2015 Creating a catalog of protein interactions. See https://doi.org/10.15363/thinklab.d85.

155. Himmelstein DS, Baranzini SE. 2016 Dhimmel/Ppi V1.0: Compiling A Human Protein Interaction Catalog. See https://doi.org/10.5281/zenodo.48443.

156. Rual J-F et al. 2005 Towards a proteome-scale map of the human protein–protein interaction network. Nature 437, 1173–1178. See https://doi.org/10.1038/nature04209.

157. Venkatesan K et al. 2008 An empirical framework for binary interactome mapping. Nat Meth 6, 83–90. See https://doi.org/10.1038/nmeth.1280.

158. Yu H et al. 2011 Next-generation sequencing to generate interactome datasets. Nat Meth 8, 478–480. See https://doi.org/10.1038/nmeth.1597.

159. Rolland T et al. 2014 A Proteome-Scale Map of the Human Interactome Network. Cell 159, 1212–1226. See https://doi.org/10.1016/j.cell.2014.10.050.

160. Menche J, Sharma A, Kitsak M, Ghiassian SD, Vidal M, Loscalzo J, Barabasi A-L. 2015 Uncovering disease-disease relationships through the incomplete interactome. Science 347, 1257601–1257601. See https://doi.org/10.1126/science.1257601.

161. Huntley RP, Sawford T, Mutowo-Meullenet P, Shypitsyna A, Bonilla C, Martin MJ, O’Donovan C. 2014 The GOA database: Gene Ontology annotation updates for 2015. Nucleic Acids Research 43, D1057–D1063. See https://doi.org/10.1093/nar/gku1113.

162. Himmelstein D, Greene C, Malladi V, Bastian F. 2015 Compiling Gene Ontology annotations into an easy-to-use format. See https://doi.org/10.15363/thinklab.d39.

163. Himmelstein D, Greene C, Malladi V, Bastian F, Baranzini S. 2015 Gene-Ontology: Initial Zenodo Release. See https://doi.org/10.5281/zenodo.21711.

164. Edgar R. 2002 Gene Expression Omnibus: NCBI gene expression and hybridization array data repository. Nucleic Acids Research 30, 207–210. See https://doi.org/10.1093/nar/30.1.207.

165. Barrett T et al. 2012 NCBI GEO: archive for functional genomics data sets–update. Nucleic Acids Research 41, D991–D995. See https://doi.org/10.1093/nar/gks1193.

166. Himmelstein D, Brueggeman L, Baranzini S. 2016 Dhimmel/Lincs V2.0: Refined Consensus Signatures From Lincs L1000. See https://doi.org/10.5281/zenodo.47223.

167. Himmelstein D, Brueggeman L, Baranzini S. 2016 l1000.db: SQLite database of LINCS L1000 metadata. See https://doi.org/10.6084/m9.figshare.3085837.v1.

168. Himmelstein D. 2016 Assessing the imputation quality of gene expression in LINCS L1000. See https://doi.org/10.15363/thinklab.d185.

169. Himmelstein D, Greene C, Jensen LJ. 2016 Positive correlations between knockdown and overexpression profiles from LINCS L1000. See https://doi.org/10.15363/thinklab.d171.

170. Himmelstein D. 2016 Announcing PharmacotherapyDB: the Open Catalog of Drug Therapies for Disease. See https://doi.org/10.15363/thinklab.d182.

171. Himmelstein D, Pouya Khankhanian, Hessler CS, Green AJ, Baranzini S. 2016 PharmacotherapyDB 1.0: the open catalog of drug therapies for disease. See https://doi.org/10.6084/m9.figshare.3103054.

172. Himmelstein DS, Pouya Khankhanian, Hessler CS, Green AJ, Baranzini SE. 2016 Dhimmel/Indications V1.0. Pharmacotherapydb: The Open Catalog Of Drug Therapies For Disease. See https://doi.org/10.5281/zenodo.47664.

173. Himmelstein D, Good B, Oprea T, McCoy A, Lizee A. 2015 How should we construct a catalog of drug indications? See https://doi.org/10.15363/thinklab.d21.

174. Wei W-Q, Cronin RM, Xu H, Lasko TA, Bastarache L, Denny JC. 2013 Development and evaluation of an ensemble resource linking medications to their indications. J Am Med Inform Assoc 20, 954–961. See https://doi.org/10.1136/amiajnl-2012-001431.

175. Khare R, Li J, Lu Z. 2014 LabeledIn: Cataloging labeled indications for human drugs. Journal of Biomedical Informatics 52, 448–456. See https://doi.org/10.1016/j.jbi.2014.08.004.

176. Khare R, Burger JD, Aberdeen JS, Tresner-Kirsch DW, Corrales TJ, Hirchman L, Lu Z. 2015 Scaling drug indication curation through crowdsourcing. Database 2015, bav016–bav016. See https://doi.org/10.1093/database/bav016.

177. Himmelstein D, Khare R. 2015 Processing LabeledIn to extract indications. See https://doi.org/10.15363/thinklab.d46.

178. McCoy AB, Wright A, Laxmisan A, Ottosen MJ, McCoy JA, Butten D, Sittig DF. 2012 Development and evaluation of a crowdsourcing methodology for knowledge base construction: identifying relationships between clinical problems and medications. J Am Med Inform Assoc 19, 713–718. See https://doi.org/10.1136/amiajnl-2012-000852.

179. Himmelstein D. 2015 Extracting indications from the ehrlink resource. See https://doi.org/10.15363/thinklab.d62.

180. Himmelstein D, Khankhanian P, Hessler C. 2015 Expert curation of our indication catalog for disease-modifying treatments. See https://doi.org/10.15363/thinklab.d95.

181. Spaulding J, Himmelstein D, Greene C, Good B. 2015 Enabling reproducibility and reuse. See https://doi.org/10.15363/thinklab.d23.

182. Hrynaszkiewicz I. 2011 The need and drive for open data in biomedical publishing. Serials: The Journal for the Serials Community 24, 31–37. See https://doi.org/10.1629/2431.

183. Molloy JC. 2011 The Open Knowledge Foundation: Open Data Means Better Science. PLoS Biol 9, e1001195. See https://doi.org/10.1371/journal.pbio.1001195.

184. McKiernan EC et al. 2016 How open science helps researchers succeed. eLife 5. See https://doi.org/10.7554/elife.16800.

185. Piwowar HA, Vision TJ. 2013 Data reuse and the open data citation advantage. PeerJ 1, e175. See https://doi.org/10.7717/peerj.175.

186. Stodden V, McNutt M, Bailey DH, Deelman E, Gil Y, Hanson B, Heroux MA, Ioannidis JPA, Taufer M. 2016 Enhancing reproducibility for computational methods. Science 354, 1240–1241. See https://doi.org/10.1126/science.aah6168.

187. Stodden V, Miguez S. 2014 Best Practices for Computational Science: Software Infrastructure and Environments for Reproducible and Extensible Research. Journal of Open Research Software 2. See https://doi.org/10.5334/jors.ay.

188. Baggerly K. 2010 Disclose all data in publications. Nature 467, 401–401. See https://doi.org/10.1038/467401b.

189. Hrynaszkiewicz I, Cockerill MJ. 2012 Open by default: a proposed copyright license and waiver agreement for open access research and data in peer-reviewed journals. BMC Research Notes 5, 494. See https://doi.org/10.1186/1756-0500-5-494.

190. Hagedorn G, Mietchen D, Morris R, Agosti D, Penev L, Berendsohn W, Hobern D. 2011 Creative Commons licenses and the non-commercial condition: Implications for the re-use of biodiversity information. ZK 150, 127–149. See https://doi.org/10.3897/zookeys.150.2189.

191. Himmelstein D, Jensen LJ. 2015 One network to rule them all. See https://doi.org/10.15363/thinklab.d102.

192. Himmelstein D, Jensen LJ, Smith M, Fortney K, Chung C. 2015 Integrating resources with disparate licensing into an open network. See https://doi.org/10.15363/thinklab.d107.

193. Oxenham S. 2016 Legal confusion threatens to slow data science. Nature 536, 16–17. See https://doi.org/10.1038/536016a.

194. Elliott R. 2005 Who owns scientific data? The impact of intellectual property rights on the scientific publication chain. Learned Publishing 18, 91–94. See https://doi.org/10.1087/0953151053584984.

195. Himmelstein D. 2015 MSigDB licensing. See https://doi.org/10.15363/thinklab.d108.

196. Himmelstein D. 2015 Incomplete Interactome licensing. See https://doi.org/10.15363/thinklab.d111.

197. Himmelstein D. 2015 LINCS L1000 licensing. See https://doi.org/10.15363/thinklab.d110.

198. Himmelstein D, Fortney K, Knox C, Southan C. 2016 Sounding the alarm on DrugBank’s new license and terms of use. See https://doi.org/10.15363/thinklab.d213.

199. Liberzon A, Subramanian A, Pinchback R, Thorvaldsdottir H, Tamayo P, Mesirov JP. 2011 Molecular signatures database (MSigDB) 3.0. Bioinformatics 27, 1739–1740. See https://doi.org/10.1093/bioinformatics/btr260.

200. Himmelstein D. 2016 Assessing the effectiveness of our hetnet permutations. See https://doi.org/10.15363/thinklab.d178.

201. Hanhijärvi S, Garriga GC, Puolamäki K. 2009 Randomization Techniques for Graphs. In Proceedings of the 2009 SIAM International Conference on Data Mining, pp. 780–791. Society for Industrial and Applied Mathematics. See https://doi.org/10.1137/1.9781611972795.67.

202. Himmelstein D. 2015 Permuting hetnets and implementing randomized edge swaps in cypher. See https://doi.org/10.15363/thinklab.d136.

203. Yoon B-H, Kim S-K, Kim S-Y. 2017 Use of Graph Database for the Integration of Heterogeneous Biological Data. Genomics Inform 15, 19. See https://doi.org/10.5808/gi.2017.15.1.19.

204. Jaiswal G. 2013 Comparative analysis of Relational and Graph databases. IOSRJEN 03, 25–27. See https://doi.org/10.9790/3021-03822527.

205. Have CT, Jensen LJ. 2013 Are graph databases ready for bioinformatics? Bioinformatics 29, 3107–3108. See https://doi.org/10.1093/bioinformatics/btt549.

206. Lysenko A, Roznovăţ IA, Saqi M, Mazein A, Rawlings CJ, Auffray C. 2016 Representing and querying disease networks using graph databases. BioData Mining 9. See https://doi.org/10.1186/s13040-016-0102-8.

207. Balaur I, Mazein A, Saqi M, Lysenko A, Rawlings CJ, Auffray C. 2016 Recon2Neo4j: applying graph database technologies for managing comprehensive genome-scale networks. Bioinformatics, btw731. See https://doi.org/10.1093/bioinformatics/btw731.

208. Summer G, Kelder T, Radonjic M, van Bilsen M, Wopereis S, Heymans S. 2016 The Network Library: a framework to rapidly integrate network biology resources. Bioinformatics 32, i473–i478. See https://doi.org/10.1093/bioinformatics/btw436.

209. Mungall CJ et al. 2016 The Monarch Initiative: an integrative data and analytic platform connecting phenotypes to genotypes across species. Nucleic Acids Res 45, D712–D722. See https://doi.org/10.1093/nar/gkw1128.

210. Himmelstein D. 2015 Using the neo4j graph database for hetnets. See https://doi.org/10.15363/thinklab.d112.

211. Himmelstein D. 2016 dhimmel/hetio v0.2.0: Neo4j export, Cypher query creation, hetnet stats, and other enhancements. See https://doi.org/10.5281/zenodo.61571.

212. Himmelstein D. 2016 Hosting Hetionet in the cloud: creating a public Neo4j instance. See https://doi.org/10.15363/thinklab.d216.

213. Belmann P, Dröge J, Bremges A, McHardy AC, Sczyrba A, Barton MD. 2015 Bioboxes: standardised containers for interchangeable bioinformatics software. GigaSci 4. See https://doi.org/10.1186/s13742-015-0087-0.

214. Beaulieu-Jones BK, Greene CS. 2017 Reproducibility of computational workflows is automated using continuous analysis. Nat Biotechnol 35, 342–346. See https://doi.org/10.1038/nbt.3780.

215. Himmelstein D, Lizee A. 2016 Estimating the complexity of hetnet traversal. See https://doi.org/10.15363/thinklab.d187.

216. Burbidge JB, Magee L, Robb AL. 1988 Alternative Transformations to Handle Extreme Values of the Dependent Variable. Journal of the American Statistical Association 83, 123–127. See https://doi.org/10.2307/2288929.

217. Himmelstein D, Khankhanian P, Lizee A. 2016 Transforming DWPCs for hetnet edge prediction. See https://doi.org/10.15363/thinklab.d193.

218. Himmelstein D. 2015 Assessing the informativeness of features. See https://doi.org/10.15363/thinklab.d115.

219. Himmelstein D. 2016 Edge dropout contamination in hetnet edge prediction. See https://doi.org/10.15363/thinklab.d215.

220. Himmelstein D. 2016 Decomposing predictions into their network support. See https://doi.org/10.15363/thinklab.d229.

221. Himmelstein D. 2016 Decomposing the DWPC to assess intermediate node or edge contributions. See https://doi.org/10.15363/thinklab.d228.

222. Lizee A, Himmelstein D. 2016 Network Edge Prediction: Estimating the prior. See https://doi.org/10.15363/thinklab.d201.

223. Lizee A, Himmelstein D. 2016 Network Edge Prediction: how to deal with self-testing. See https://doi.org/10.15363/thinklab.d194.

224. Himmelstein D. 2016 Cataloging drug–disease therapies in the ClinicalTrials.gov database. See https://doi.org/10.15363/thinklab.d212.

225. Brown AS, Patel CJ. 2017 A standard database for drug repositioning. Sci. Data 4, 170029. See https://doi.org/10.1038/sdata.2017.29.

226. Shameer K et al. 2017 Systematic analyses of drugs and disease indications in RepurposeDB reveal pharmacological, biological and epidemiological factors influencing drug repositioning. Briefings in Bioinformatics See https://doi.org/10.1093/bib/bbw136.

227. Sharp ME. 2017 Toward a comprehensive drug ontology: extraction of drug-indication relations from diverse information sources. J Biomed Semant 8. See https://doi.org/10.1186/s13326-016-0110-0.

228. Himmelstein D, Lizee A, Hessler C, Brueggeman L, Chen S, Hadley D, Green A, Khankhanian P, Baranzini S. 2015 Rephetio: Repurposing drugs on a hetnet [proposal]. See https://doi.org/10.15363/thinklab.a5.

229. Himmelstein D, Lizee A. 2016 Measuring user contribution and content creation. See https://doi.org/10.15363/thinklab.d200.

230. Patil C, Siegel V. 2009 This revolution will be digitized: online tools for radical collaboration. Disease Models & Mechanisms 2, 201–205. See https://doi.org/10.1242/dmm.003285.

231. Mietchen D, Mounce R, Penev L. 2015 Publishing the research process. RIO 1, e7547. See https://doi.org/10.3897/rio.1.e7547.

232. Powell K. 2016 Does it take too long to publish research? Nature 530, 148–151. See https://doi.org/10.1038/530148a.

233. Vale RD. 2015 Accelerating scientific publication in biology. Proc Natl Acad Sci USA 112, 13439–13446. See https://doi.org/10.1073/pnas.1511912112.

234. Allison DB, Brown AW, George BJ, Kaiser KA. 2016 Reproducibility: A tragedy of errors. Nature 530, 27–29. See https://doi.org/10.1038/530027a.

235. Himmelstein D et al. 2016 Workshop to analyze LINCS data for the Systems Pharmacology course at UCSF. See https://doi.org/10.15363/thinklab.d181.

236. Waldrop MM. 2015 Why we are teaching science wrong, and how to make it right. Nature 523, 272–274. See https://doi.org/10.1038/523272a.

237. Giles J. 2012 Going paperless: The digital lab. Nature 481, 430–431. See https://doi.org/10.1038/481430a.

